# How host contact network impacts *N*-strain SIS dynamics with coinfection via a global replicator equation

**DOI:** 10.64898/2026.01.23.701350

**Authors:** Nicola Cinardi, Sten Madec, Erida Gjini

## Abstract

Although the effects of contact network heterogeneity on epidemic spreading are well established, its influence on multi-strain selection and coexistence in endemic scenarios remains comparatively understudied. Here, we develop an SIS framework for interacting strains with co-infection, which, under similarity, reduces to replicator dynamics on a host contact network. Leveraging its analytical tractability, we investigate how network heterogeneity shapes multi-strain dynamics relative to homogeneous contact distributions. While the model accounts for variation across five epidemiologically relevant traits, we focus on strains differing only in their co-infection vulnerability matrix. We show that network heterogeneity modifies pairwise invasion fitness, affecting both the timescales and outcomes of multi-strain dynamics. Increasing heterogeneity generally accelerates strain dynamics and reduces the number of coexisting strains. The magnitude of fitness variations due to network structure is more pronounced at low transmission intensity and low co-infection vulnerability (low *R*_0_ → 1 and *σ*→ 0), but these regimes imply also lower system’s intrinsic susceptibility to heterogeneity. For large *N*, greater antisymmetry in the homogeneous replicator matrix at low *R*_0_ and *σ* implies lower susceptibility but larger heterogeneity-induced effects. Conversely, weaker antisymmetry at higher *R*_0_ and *σ* implies greater susceptibility but smaller effects. As a consequence, effective dynamical sensitivity to heterogeneity is expected to peak at intermediate transmission intensity and co-infection facilitation. Despite the complexity of replicator dynamics for high number of strains, this framework reduces dimensionality of the original epidemiological system, enabling efficient simulations and direct quantification of network effects. Its bottom-up parameterization further provides insight into epidemiological transitions across macroscopic gradients and may guide interventions against polymorphic infectious agents.

## 1 Introduction

The spread of infectious agents in populations is a fundamental problem with profound implications for public health, ecology, and socio-cultural systems. Traditionally modeled through compartmental frameworks such as the SIR (*Susceptible-Infected-Recovered* ) or SIS (*Susceptible-Infected-Susceptible*) models, infection/contagion dynamics is approximated by mass-action assumptions for contacts and transmission, but it is currently widely recognized that they are significantly influenced by the structure of the underlying host population and heterogeneity in their contact patterns [25, 23], infectivity or susceptibility to disease [13].

In recent years, complex network theory has emerged as a powerful framework for capturing the heterogeneous topologies characteristic of real-world contact patterns ranging from social interactions to infrastructure and biological networks [3, 2]. The study of epidemic processes on host contact networks, under varying topologies, has revealed rich dynamical behaviors, such as new threshold phenomena, localization, and persistence [10, 24, 29, 39, 27].

Similarly the study of SIS dynamics on host contact networks has received attention for how network features, i.e. degree variance or topology, impact the endemic steady state [2]. Yet, the study of multi-strain infection processes on host contact networks has been more limited and only restricted to special cases [15, 6, 34].

It is widely appreciated that pathogen populations are heterogeneous, and that the dynamics of infectious diseases are often far more intricate than those captured by classical single-pathogen models. In many real-world scenarios, multiple strains or variants of a pathogen co-circulate, each with distinct transmission characteristics, immune responses, or treatment outcomes - traits via which they may interact at the within-host or between-host level.

Of course introducing such diversity into epidemiological models makes them high-dimensional and more complex. And typically analysis is restricted to a few strains, or only special cases with *N* strains. Multi-strain epidemic dynamics on host contact networks has also obeyed the same trend, with studies mainly focusing on *N* = 2 or *N* = 3 strain scenarios [5, 15, 34], and typically adopting computational simulations for examination of the dynamics. For example [15] investigate a SIS dynamics with two diseases/strains comparing high-clustered versus random networks (a network with the same degree distribution as the high-clustered one but with randomly rewired links). They find that a microscopic increase in transmissibility can cause a macroscopic difference in the expected epidemic size on a clustered network but not on an otherwise equivalent random network. In [34] the authors study a SIS model with 3 diseases (2 pathogens of which one in two strains). They only study the Erdos–Rényi graph (and compare it with a homogeneously mixed population). They study competition and cooperation together and find that cooperation causes discontinuous transitions where the probability of an outbreak and prevalence change abruptly around a critical value of the transmission rate. This phenomenon, however, is sensitive to the network topology.

Despite the insights gained by previous studies, a general understanding for how *N* -strain dynamics in endemic SIS scenarios with coinfection unfold on a host contact network is still missing. In this work, we aim to fill this gap, bridging between the complexity of *N* strain epidemiological models and host contact networks. We address especially the important, and less well-studied topic of coinfection on host contact networks, where *N* strains co-circulate in a SIS fashion and may interact via several fitness dimensions. Such multi-strain and co-infection scenarios may be relevant for viral transmission (e.g., influenza, HIV, SARS-Cov2), polymorphic bacteria colonization and co-colonization processes (e.g. *Streptococcus pneumoniae*), and even more generally information spreading in social systems. Understanding the interaction and selection processes among multiple infectious agents is crucial for predicting their dynamics and designing effective control strategies.

We have extended a previous framework for modeling SIS dynamics with coinfection and *N* interacting strains, based on strain similarity and host homogeneous mixing [22, 18]. In [21] we have studied the same system but on a host contact network and we have shown, like under the homogeneous mixing assumption, that in the host contact network case, a global replicator equation emerges for global selection dynamics between strains on the network. The parameters of this replicator contain the trace of the relevant heterogeneity in the network and of the relevant trait asymmetries between strains. Here we will study this particular replicator, responsible for strain dynamics on the network and examine the effect of network characteristics, topology and other global parameters, on the sensitivity and quality of strain selection dynamics. It is not immediately clear that in a structured population, similar to the homogeneous mixing case, dynamics with interacting strains leads to the replicator equation. Even less obvious is how the structure of the network impacts such dynamics and which outcome will be favored given certain initial conditions and macroscopic parameters. No previous work has addressed these two problems. In [21] we detailed the theoretical approach to address the first, in this work we focus on the answer to the second, by showing the role of properties of population contact networks on multi-strain interactions. The paper is organized as follows: in Section 2 we present the general coinfection model on a host contact network and its extension to multiple interacting strains, providing the link with the replicator dynamics. In Section 3 we explore global epidemiological variables as a function of network parameters, and in Section 4 we investigate several metrics of multi-strain dynamics sensitivity to host contact network heterogeneity, both in the *N* = 2 and larger *N* setting. We conclude with a discussion and outlook for the future.

## 2 General coinfection model on a host contact network

We start by considering an SIS model with coinfection (SID), on a host network without explicitly distinguishing strain identities. Each strain has transmission probability *β* per contact^1^, from an infected host, and each infection episode lasts on average *γ*^−1^ units of time. Singly-infected hosts can be coinfected by acquiring a second infection of the same or a different strain. All the infections are indistinguishable epidemiologically, except for the transition from single to co-infection, which happens with an altered coefficient *σ*, relative to the transition from susceptible to single-infected. This coefficient can be below or above 1, indicating inhibition or facilitation effects between primary colonization and co-colonization respectively.

This serves as a starting point to obtain general properties of the endemic equilibrium for an SIS model with coinfection (SID). The assumptions are similar to those appearing in previous papers, under the homogeneous mixing hypothesis [12, 22].

The difference is that in the current formulation, the host population is structured as a network, thus we follow in spirit the SIS mean-field approaches delineated for network models [30, 31]. Each individual (node of the network) has a potentially different number of contacts, given by *k. P* (*k*) is the degree distribution of the network giving the probability of finding a node of degree *k* within the network. Here we use the following notation when referring to a particular network *N* = {*K, P*}, where K = {*k*_1_, … *k*_*L*_} ⊂ ℕrepresents all possible degrees of the nodes and *P* = (*p*_*k*_)_*k*∈*K*_ represents the probability distribution of such degrees. Several prototype networks exist and have been studied in the literature with specific signatures on epidemic dynamics and thresholds (Table 1).

**Table 1:**
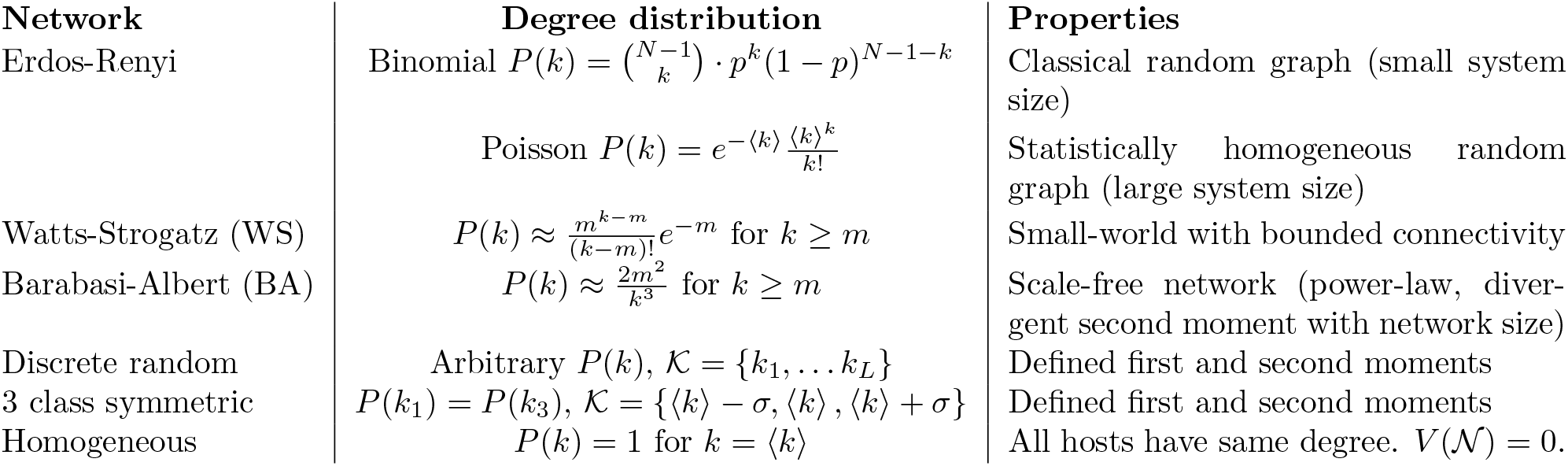
Prototype connectivity structures that can be studied for epidemic processes on explicit host contact networks. The second and third distributions are general models that lead to networks with the referred properties but do not constitute exact expressions. The last three distributions are the ones selected for the simulations in this paper, due to their simplicity and generality, allowing for conclusions that do not depend on any specific topology. Our framework presented here is general and can be applied to other finite degree distributions.

| **Network** | **Degree distribution** | **Properties** |
|---|---|---|
| Erdos-Renyi | Binomial $P(k) = \binom{N-1}{k} \cdot p^k(1-p)^{N-1-k}$ | Classical random graph (small system size) |
|  | Poisson $P(k) = e^{-\langle k \rangle} \frac{\langle k \rangle^k}{k!}$ | Statistically homogeneous random graph (large system size) |
| Watts-Strogatz (WS) | $P(k) \approx \frac{m^{k-m}}{(k-m)!}e^{-m}$ for $k \geq m$ | Small-world with bounded connectivity |
| Barabasi-Albert (BA) | $P(k) \approx \frac{2m^2}{k^3}$ for $k \geq m$ | Scale-free network (power-law, diver-gent second moment with network size) |
| Discrete random | Arbitrary $P(k)$, $\mathcal{K} = \{k_1, \dots k_L\}$ | Defined first and second moments |
| 3 class symmetric | $P(k_1) = P(k_3)$, $\mathcal{K} = \{\langle k \rangle - \sigma, \langle k \rangle, \langle k \rangle + \sigma\}$ | Defined first and second moments |
| Homogeneous | $P(k) = 1$ for $k = \langle k \rangle$ | All hosts have same degree. $V(\mathcal{N}) = 0$. |

Remember that ⟨*k*⟩ = Σ_*k*∈*K*_ *kp*_*k*_ is the mean degree and that the second moment of the degree distribution is *k*^2^ = Σ _*k*∈*K*_ *k*^2^*P*_*k*_; we denote *V* (*N*) = ( *k*^2^ −⟨*k*⟩^2^) the variance of the degree distribution^2^.

Hence, we consider the hosts as structured in discrete homogeneous subclasses, defined in this case by contact number

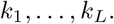

Host demographic rates are given by *r*_*k*_ (class-specific birth) and *r* (death) for each contact (degree) class then the total number of hosts in each class is given by 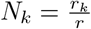 and the preserved contact degree distribution is

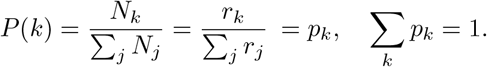

The subdivision into degree classes is done for analytical convenience; each class is connected to members of the other classes, and all together form the full network.

In [21], along the same similarity arguments and singular perturbation approach of [22, 18], we derived in detail how epidemiological SIS dynamics involving multiple strains and coinfection on a host contact network (away from the homogeneous mixing assumption) gives rise to a global replicator dynamics.

The replicator equation is a central framework in evolutionary game theory that models the frequency-dependent success of competing strategies [14], and in this kind of model, appearing under a slow-fast framework, like in previous studies [22, 18], it has twofold applications. First, it helps to study multistrain selection in a compact, elegant and analytically-tracatable manner and, secondly, it reduces the original model dimensionality therefore enabling faster and more efficient computations for arbitrary strain number.

Specifically in the theoretical paper [21], we demonstrated that under plausible assumptions (e.g., similarity of strains), the evolution of strain dynamics can be effectively captured by a replicator equation. The explicit details of this mapping, from epidemiology on the network to the replicator equation, reveal how strain frequencies evolve not merely due to their intrinsic trait asymmetries and pairwise selection advantages, but also due to the properties of connectivity of hosts in the network.

In the following two sections we describe briefly the general framework, and subsequently we focus on our main question in the current paper: What is the impact of the network heterogeneity on the emerging replicator dynamics and ultimately on strain selection?

### 2.1 Deterministic SID model for degree classes

We consider a structured transmission model, based on [22], where the transmission dynamics are specified in each contact class *k* (denoting the compartment of hosts with *k* contacts per unit of time). We will model the rates of change in susceptible *S*_*k*_, single-infected *I*_*k*_ and co-infected hosts *D*_*k*_ within each such degree class, assuming that all nodes (individuals) with the same degree are statistically equivalent.

Since the number of hosts in each class is the fixed quantity 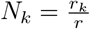, we renormalise each class by dividing by *N*_*k*_. Thus, the per-capita birth and death rates in each class are given by the common value *r* = *r*_*k*_*/N*_*k*_. The model variables after rescaling, *S*_*k*_, *I*_*k*_, *D*_*k*_, ∀ *k* ∈ *K* represent the density of Susceptible, single-Infected, and Double-infected hosts among all hosts who have *k* connections. Their dynamics satisfies

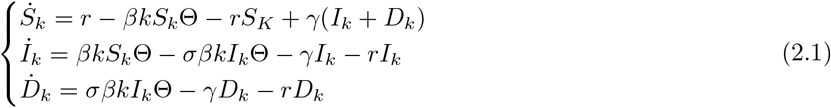

where *β* represents the probability of infection per contact. In the model *γ* is the clearance rate for both single- and co-infection, while *σ* is the relative vulnerability to co-infection when a host is singly-infected. When *σ <* 1 it describes antagonistic (competition) interaction between single and coinfection. Conversely, when *σ* ≥1 it describes facilitative (cooperation) interaction between single and coinfection. In the network, under lack of degree correlations, within each connectivity class, infection happens according to a global force of infection, namely the mean-field parameter dictating the probability to be in contact with an infected node (which is independent of *k*):

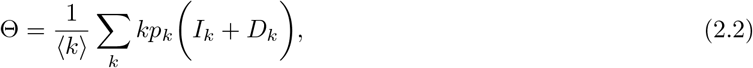

where ⟨*k*⟩ is the average degree and *p*_*k*_ is the network degree probability distribution. Notice that Θ accounts for contributions by all single- and double-infected nodes in the network and connects all the degree classes. In fact, it gives the probability to get in contact with an infected host over the whole network, similar to previously-studied SIS models on networks [31].

An essential assumption of the framework is that the population density remains fixed within each degree class. This allows us to work with normalised variables satisfying *S*_*k*_ + *I*_*k*_ + *D*_*k*_ = 1 for every *k*.

Under this assumption, and denoting *T*_*k*_ = 1 −*S*_*k*_ = *I*_*k*_ + *D*_*k*_, the dynamics is essentially driven by

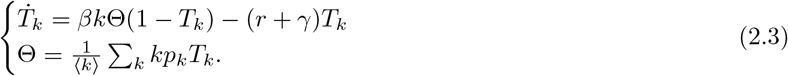

which is a well studied SIS model on network ([20, 40]). The other equations on *I*_*k*_ and *D*_*k*_ are to close the analysis of this SID system on a network.

Shortly, if (2.3) has a positive steady state, then Θ is a solution of

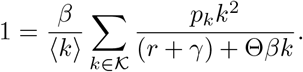

In [21], and consistent with the epidemic thresholds known for classical SIR and SIS models on networks (see [31, 25]), we show that this equation has a unique solution Θ^*^ ∈ (0, 1) if and only if 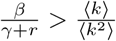.

Once Θ^*^ is determined, we recover explicitly the asymptotic proportions of susceptible, total infected, single-infected and co-infected hosts in each degree class:

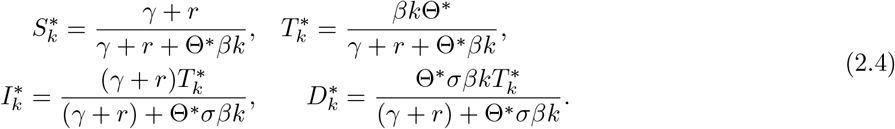

Written in terms of class-specific reproduction numbers *R*_*k*_ = *βk/*(*γ* + *r*), this endemic equilibrium becomes:]

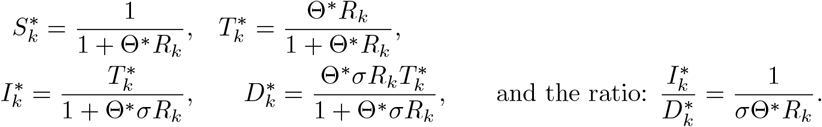

Finally, this endemic equilibrium is globally stable whenever it exists.

Written in terms of a super-critical net reproduction number for the network, this implies that (2.1) has a global endemic attractor if an only if

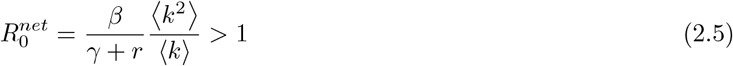

which defines the epidemic threshold condition as:

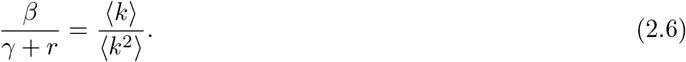

Such threshold invokes a key relationship between the first and second moment of the host contact degree distribution, and illustrates that in networks displaying a larger heterogeneity ratio, ⟨*k*^2^*⟩/⟨ k⟩* the epidemic threshold is reduced.

### 2.2 Multi-strain co-infection model extension

What if instead of a single strain we have *multiple* strains that can co-circulate and co-infect hosts in a population? These strains can be different in their transmission (*β*^*i*^) and clearance traits (*γ*^*i*^, *γ*^*ij*^), in their co-infection vulnerabilities (*σ* ^*ij*^) manifested as pairwise competition or cooperation (denoted as *k*_*ij*_ in [22, 18]), and in their priority effects for transmission from co-infection. To accommodate such diversity resolution between strains, we update the model. Let us denote the host degree class with a subscript *k* and the colonization status by different strains in the superscripts (*i* for single infection, and *ij* for co-infection). In principle strains can be different along 5 fitness dimensions [18], including transmission probabilities from single and coinfected hosts, duration of single and coinfection, and coinfection vulnerabilities (see §2.2.1for parameter details).

We now have two layers of structure intertwined in our compartmental model variables (see Figure 1), for *contact number* and *strain*:

**Figure 1:**
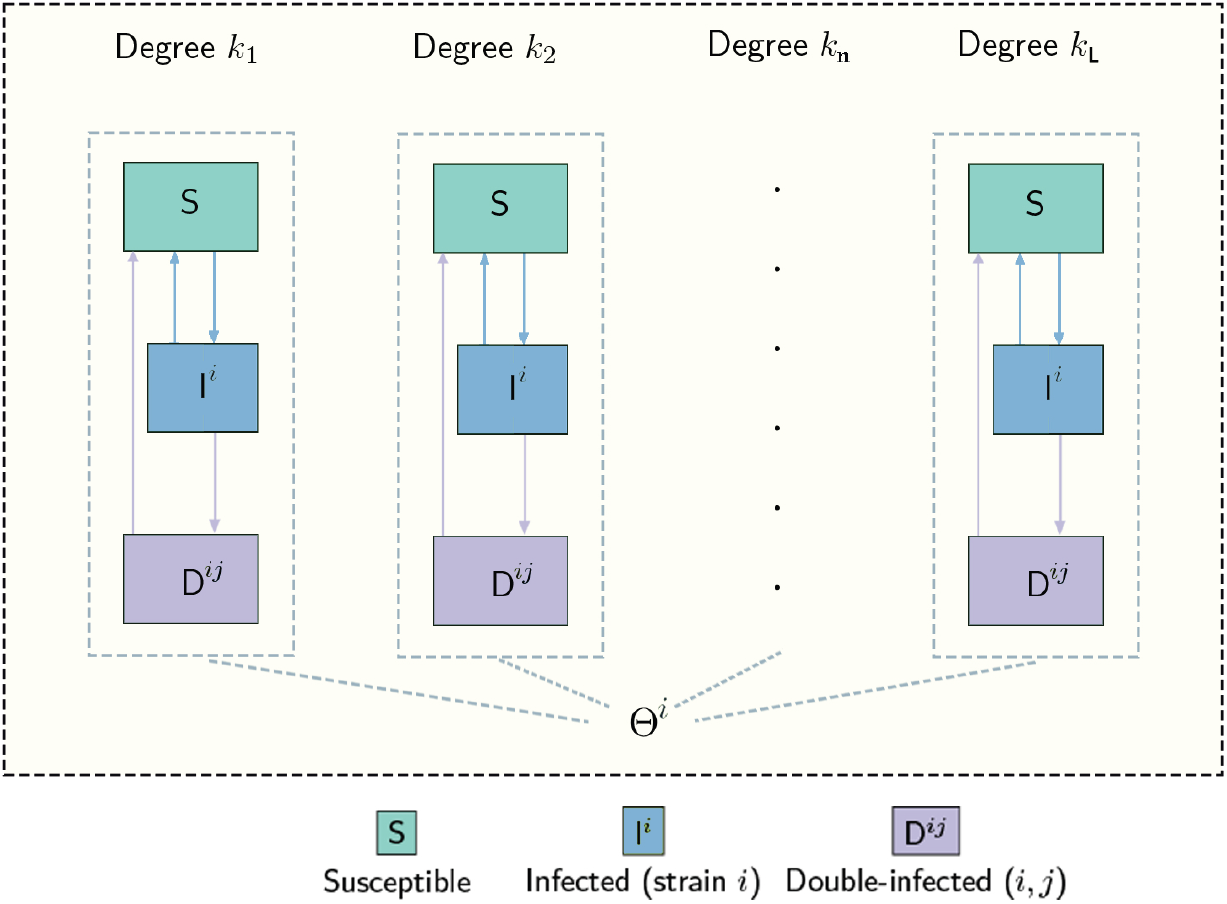
Graphical representation of host population structure by contact degree-class and epidemiological infection status. In each degree class, we find Susceptibles (*S*), Single-Infected hosts by strain *i*, (*i* = 1, ..*N* ), and Double-Infected hosts by strains (*i, j*) including double-infection by the same strain. Each individual is part of a given class *k* as it is connected to *k* other hosts. The statistical equivalence assumption between hosts of the same degree allows an analytical treatment through the mean-field approach. All the classes are connected to form the full network, and the interactions are described through the mean-field emergent quantity Θ which gives the probability to get in contact with an infected host (Θ^*i*^ referring to hosts that can transmit strain *i*).

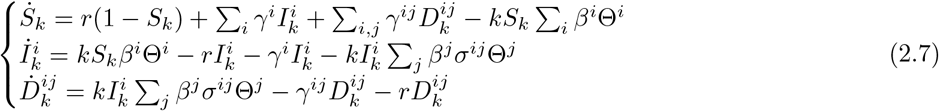

with a strain-specific global probability to be in contact with a node transmitting strain *i* given by:

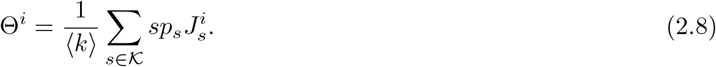

We may alternatively write 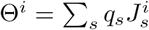, by posing 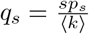, which represents the probability to encounter a host from (degree-)class *s*. From now on, we use *s* when summing over the degrees *K* = {*k*_1_, …, *k*_*L*_ *}*. In the equations here we assume no bias in transmission from mixed coinfection, making each of the two strains equally probable to be transmitted from *D*^*ij*^, (*i* ≠ *j*) hosts. Thus the density of hosts of class *s* that can transmit strain *i* is in general given by:

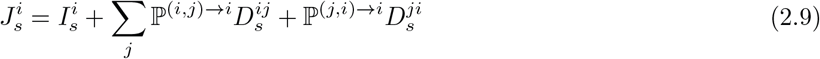

which, in the special case of no priority effects in transmission from coinfected hosts harbouring different strains 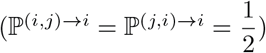, becomes simply:

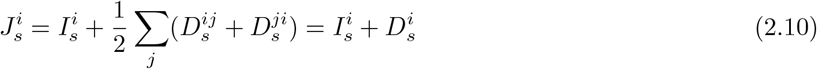

under the definition 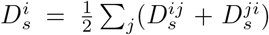, accounting for probability equal to 1/2 of each strain to be transmitted from a mixed coinfection host, as in [22].

#### 2.2.1 Under strain similarity, the replicator equation emerges

The traits allowed to vary between any two strains in this model are: i) *transmission probability β*^*i*^, ii) *duration of single-* and iii) *co-infection* via clearance rates *γ*^*i*^,*γ*^*ij*^, iv) *transmission bias from coinfection via order of arrival ℙ* ^(*i,j*)→*i*^, and v) pairwise *co-infection vulnerabilities σ*^*ij*^, as specified also in [18].

When we study the quasi-neutral formulation of this model, similar to [18], for which all traits between strains can be assumed to be relatively similar, and expressed as a reference value plus a small deviation from that value, we effectively describe the traits, using the small parameter 0 *< ϵ* ≪ 1, as follows:

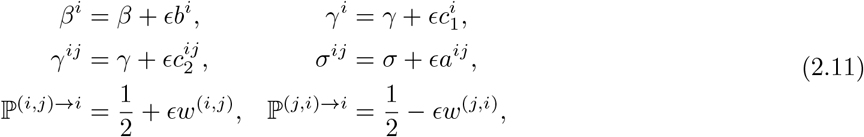

and seek a model reduction into fast and slow dynamics to simplify analysis and computation. It is precisely this assumption 2.11 that is used in our theoretical paper [21] to derive a very general slow-fast approximation of the overall epidemiological dynamics on the network. Being outside the scope of this paper, here we only briefly outline the strategy that we adopt in [21] to arrive at the replicator equation, and present the key ingredients that allow us to focus, in this work, on the impact of the network and the resulting network-dependent replicator system. The key assumption is the quasi-neutrality, which allows us to use Tikhonov’s theorem after rewriting the system in a slow–fast form, in which the fast dynamics are completely determined by the global variables, and a linear system asymptotically governed by a matrix *A*^*^ whose dominant eigenvalue is equal to 0. With this, it is possible to derive the reduced slow dynamics, which takes the form of a replicator equation. It is therefore required to perform the spectrum analysis of a large matrix. To overcome this difficulty, one can rewrite the system such that *A*^*^ has a Metzler structure, hence allowing a detailed spectral analysis. In what follows, we take the slow-fast approximation as a starting point, and focus on the general implications of the reduced model. We discuss in detail the key ingredients in the following paragraphs.

##### Fast variables

The fast variables in the system are the proportions of susceptible, single-infected and coinfected hosts in each connectivity class, tending to a steady-state (implicit) endemic equilibrium, given by Eqs. 2.4:

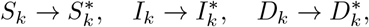

with 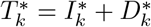 the density of total infected hosts of degree *k* at the steady state.

##### Slow variables

Like we show in the theoretical paper [21], the slow variables in the system are the *N* strain frequencies throughout the entire network, satisfying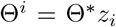. They are governed by a replicator equation over the whole network for *τ >* 0:

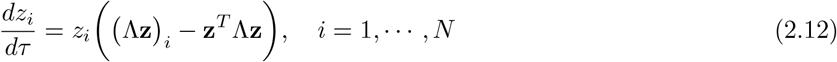

where Λ, the global pairwise fitness matrix between strains, is explicitly related to the contact-degree distribution among hosts, as well as various trait asymmetries between strains (see §2.2.2).

#### 2.2.2 Global pairwise invasion fitness matrix between strains in a host contact network

The matrix appearing in the global replicator equation (2.12) can be written in terms of components, which are a suitably-weighted average over perturbations in 5 fitness dimensions between strains (5 traits):

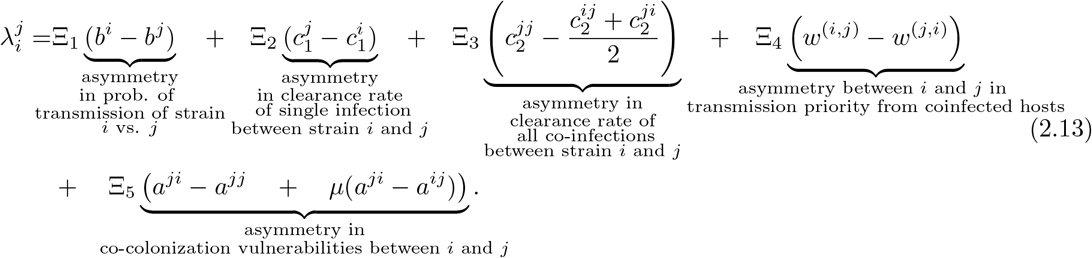

The traits varying between any two strains are: i) *transmission probability β*^*i*^, ii) *duration of single-* and iii) *co-infection* via clearance rates *γ*^*i*^,*γ*^*ij*^, iv) *transmission bias from coinfection via order of arrival ℙ* ^(*i,j*)→*i*^, and v) *co-colonization vulnerabilities σ*^*ij*^ [18], as specified in (2.11). The weights for each trait Ξ_*i*_ *>* 0, scaling their contribution to pairwise invasion fitness 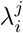, depend on the emergent network mean-field quantities Ξ_*k*_ which are given in the Supplementary section S1, Eq. (S1.1), and on the mean ratio *µ* of single- to co-infection in the system which is obtained as a nontrival weighted average.

##### Class-specific reproduction numbers

For each contact degree, a class-specific reproduction number emerges and is given by

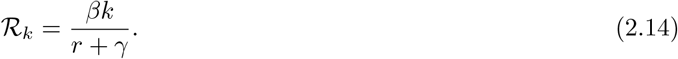

##### Ratio of single-to-co-infection

In the network model, we first have a ratio of single-to-coinfection in each degree class:

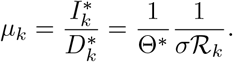

We also have the global mean ratio of single- to co-infection in the system which is not *I*^*^*/D*^*^, but is instead given by a nontrivial weighted average of the ratios *µ*_*k*_ in each degree class:

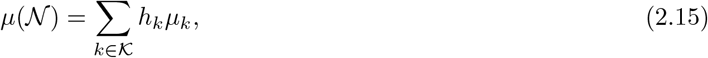

with 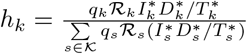, where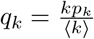 . In these weights, global class-specific aggregated quantities appear and interact nonlinearly with each other to shape the importance of each trait axis on final relative fitnesses between strains.

##### Special case: Strain variation only in co-colonization susceptibilities

It is important to notice, that unlike the analytical transparency of the unstructured model [22], a general analytical expression of the Λ matrix between strains corresponding to a host contact network is not attainable (for instance, an explicit formulation of *µ* is not always possible), but a numerical investigation for specific cases can be performed (see discussion in [21]). Such one special case will be the focus of our investigations in the remaining sections.

In the case we focus on for illustration, strains are equal along all fitness dimensions, except for pairwise vulnerabilities to co-colonization *σ* ^*ij*^ = *σ* + *ϵa*^*ij*^, (*ϵ* small) and the pairwise invasion fitness between any two strains over the entire network (Eq.2.13) takes a simpler form, given by:

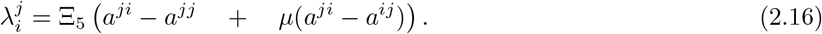

This case is sufficiently general to generate all types of replicator dynamics and has the peculiar feature of Λ dependence on *µ* - shown to act as a contextual tuning parameter for the quality of coexistence dynamics in the homogeneous mixing case [11].

### 2.3 Closure of the model reduction

Combining Eqs. (2.4) and (2.12), the infection compartments of the full network for each class *k* and epidemiological status, are obtained as:

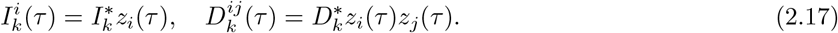

#### Global epidemiological variables

Finally, from the proportions of susceptible, single-infected, and co-infected hosts in each connectivity class, we recover the global variables of the epidemiological model as:

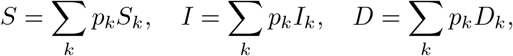

which, for 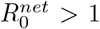, tend to their endemic steady state values given by the sum over the degree classes (Eq. 2.4)

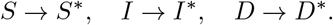

Of course, the actual dynamics in this slow-fast formulation is augmented with appropriate initial conditions, **z** such that **z**(0) = **z**^0^. If all the trait perturbations between strains do not depend on the class *k*, but only on the strains, like in 2.11, the replicator equation has a form which is very similar to the replicator equation for only one class of hosts (see [18]).

## 3 The network effect on global epidemiological quantities

### Θ^*^ and global endemic prevalence

We first study how global epidemiological quantities: Θ^*^, 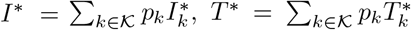 vary as a function of network heterogeneity. In Figure 2 we illustrate how they vary with the coefficient of variation (CV) of the degree distribution,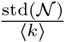. In these numerical simulations, we observe that Θ^*^ always increases with network variability. For too low mean degree, ⟨*k*⟩ = 8, we have sub-critical 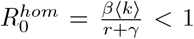 and the disease is not able to persist in relatively more homogeneous networks (i.e. small *std*(*N*)). This yields the typical bifurcation behavior in networks expected at 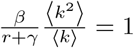. For the other values of ⟨*k*⟩ the disease persists endemically for any *std*(*N*). In Fig.2b-d, ⟨ *k*⟩ is fixed at the values 11, 15, and 30 respectively, and we plot the total prevalence of infection *T* ^*^ which differently from Θ^*^ can increase or decrease with network variability, and does not depend on *σ* . In these plots, the behavior *I*^*^ is also depicted for two values of *σ* . Note that *I*^*^ + *D*^*^ = *T* ^*^ so the gap between *I*^*^ and *T* ^*^ gives exactly the prevalence of the co-infected hosts *D*^*^.

**Figure 2:**
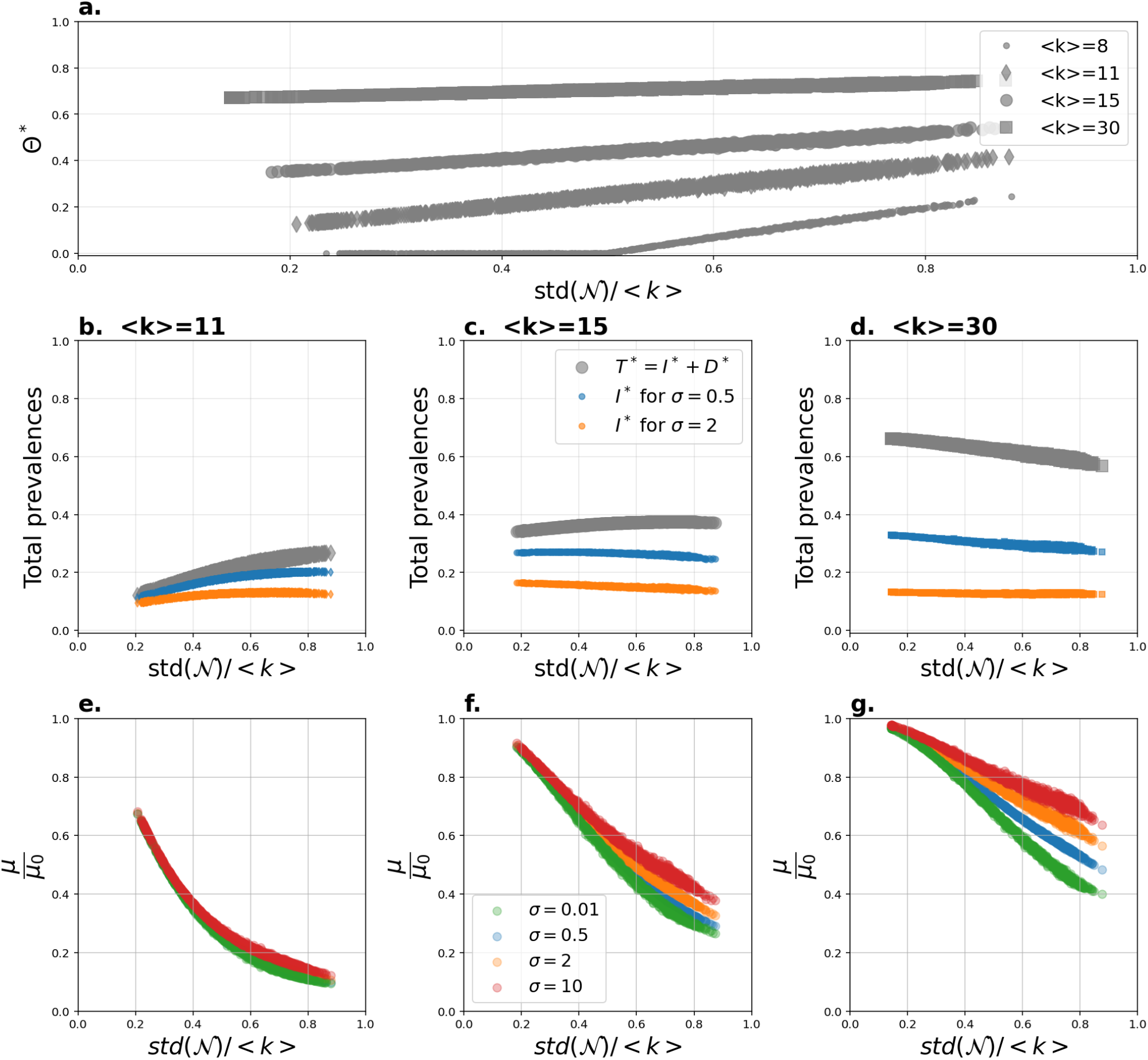
Global epidemiological quantities of the multi-strain SIS model with coinfection on host contact networks, as a function of network heterogeneity. We illustrate how global epidemiological quantities: Θ^*^, *I*^*^, *T* ^*^ and *µ*(*N*) vary as a function of network heterogeneity, here modulated via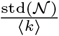. In these numerical simulations, we set 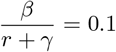. For each fixed mean degree *k* and co-infection susceptibility factor *σ* we compute these quantities from 1000 random networks generated without any specific topology. **a**. We plot the global probability of infection Θ^*^ for 4 values of ⟨*k*⟩ ∈ {8, 11, 15, 30}. **b. c. d**. Here ⟨*k*⟩ is fixed at the values 11, 15, and 30 respectively, and we plot the total prevalence of infection *T*^*^ which does not depend on *σ*, and *I*^*^ for two values of *σ* . Note that *I*^*^ + *D*^*^ = *T*^*^ so the gap between *I*^*^ and *T* ^*^ gives exactly the prevalence of the co-infected hosts *D*^*^. **e. f. g**. Here, for the same three fixed values of ⟨*k*⟩, we plot the ratio *µ*(N)*/µ*(*N*_0_) for different values of the co-infection parameter *σ* (for brevity written as *µ/µ*_0_) and high values of this ratio favouring complex attractors [11]. For the simulations that we will perform on the heterogeneous network, we will also explore such tuning effects, focusing on the pairwise invasion fitness matrix as a function of global aggregate quantities.

### The effect of single- to co-infection ratio

*µ* As shown in Section 2.2.2, in the network model, the mean ratio of single- to co-infection in the system is more complicated (Eq. 2.15), given by a nontrivial weighted average of the ratio 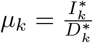 in each degree class: *µ*(*N*).

The dependence of this quantity on the coefficient of variation of the network is illustrated in Fig2 eg, confirming that it decreases with network variability, and that it is higher for higher strain cooperation parameter *σ*, and for higher transmission intensity (here mean contact degree) ⟨ *k*⟩ .

Relative to the homogeneous case, for the network we can prove using Jensen’s Inequality (see our Theoretical paper [21]) that for a fixed average degree ⟨ *k*⟩, the global ratio of single to coinfection in a heterogeneous network is always lower than the corresponding ratio in a homogeneous population with equal mean:

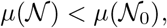

where *N, N*_0_ indicate the general heterogeneous network and the homogeneous one with same mean degree. This feature is also confirmed in our simulations in Fig.2 e-g, where besides the decreasing relationship with the coefficient of variation, the ratio *µ*(*N* )*/µ*(*N*_0_) is always below 1.

The significance of this result for multi-strain dynamics will be considered in the next section. Indeed, in what concerns Λ, we allow variations only in the component of the pairwise fitness matrix coming from co-colonization susceptibility factors within- and between-strains (Eq. 2.16), assuming all the others are equal to zero. Consequently, the effects of *µ*(*N*) on the multi-strain dynamics on the network will come only from these perturbations (see (§2.2.2)). Thus, from the relative comparison *µ*(*N*) *< µ*(*N*_0_) and from the decreasing relationship with network degree heterogeneity, in light of our previous findings [11], keeping all else fixed, we hypothesize naturally that the effect of first, the network itself, and secondly, of network heterogeneity, on strain dynamics may be a stabilizing one, inducing shifts like those expected in the *µ*→ 0 limit in the unstructured population case [11]. Similarly, for fixed coefficient of variation, as the interaction among strains in coinfection (*σ* ) tends towards competition (values below 1 approaching 0), *µ*(*N* ) also gets lower, suggesting again multi-strain stabilization effects like those found in [11].

## 4 The impact of host contact network on multi-strain dynamics

Although this replicator equation (Eq. 2.12) is general and allows to study multiple trait variations between strains simultaneously (see §2.2.2), in this work, we focus only on variations on the perturbation terms related to the co-colonization vulnerabilities between strains, like in [22, 11]. In this scenario, it was shown for a homogeneous population that the ratio of single infection to co-infection *µ* = *I/D* in the system, acts as a modifier of dynamic complexity between strains, with low values of this ratio favouring multi-stable coexistence

Thus, in this section, through simulations, we ask the question of what the difference is between a “homogeneous” network and a “heterogeneous” one with regards to multi-strain SIS dynamics and coinfection. To what extent does network heterogeneity or other epidemiological parameters have an impact? We will consider mean-field parameters as relevant ‘bifurcation’ or tuning parameters for the selection dynamics between strains. Thus, we will study the effect of host contact network on strain dynamics, assuming the strains share the same transmission and clearance rates, but they differ in the pairwise susceptibilities to co-colonization. These pairwise susceptibilities are given by 4 coefficients for each pair of strains: within- and between-strain coinfection. We will allow strains to differ from neutrality *σ* ^*ij*^ = *σ* + *εa*^*ij*^ in eq. (2.11) only in co-colonization coefficients, hence the matrix *A* = (*a*^*ij*^)_1≤*i,j*≤2_ captures such standardized interaction asymmetries, whereas *σ* captures mean-field interaction.

Each host contact network that we will study is denoted by *N* = {*K, P*}, where *P* = (*p*_*k*_)_*k*∈*K*_ represents the degree distribution with mean *k* and variance *V* (*N*) = (⟨*k*^2^⟩ −⟨*k*⟩ ^2^). We will consider networks that vary in mean and/or variance of degree distribution, as well as the functional form of the degree distribution. In our simulations, we will often refer to the standard deviation of the host degree distribution *std*(*N*), and to the basic reproduction number of the corresponding homogeneous network with all hosts having identical degree equal to the mean degree ⟨*k*⟩ as *R*_0_.^3^.

So the dimensions of variation we will consider are the following:

- Mean transmission intensity, as captured by *R*_0_ of the strictly homogeneous degree model with same ⟨*k*⟩
- Mean susceptibility to coinfection, as captured by *σ*
- Asymmetry structure of pairwise coinfection susceptibilities, *A* matrix
- Standard deviation of the degree distribution keeping the mean ⟨*k*⟩ fixed.

### 4.1 Impact of network heterogeneity - the case of *N* = 2 **strains**

Initially we fix here all the epidemiological parameters and restrict our attention to a system of only 2 strains. To understand how a host contact network *N* influences the strain dynamics, we first define the reference homogeneous network *N*_0_ ={⟨*k*⟩, 1} - a network with only one degree class, in which all the nodes have the same degree - and we fix the rescaled interaction matrix *A*.

#### The role of network variance and strain interaction asymmetry on pairwise invasion fitness

From the exposition in the previous section, we know that the replicator equation for the strain dynamics over the network critically depends on the pairwise fitness matrix 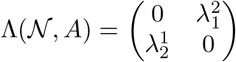 whose components we write as 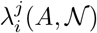. Thus, the natural way to compare the effect of the network is to compare how the invasion fitness matrix changes, Λ(*A, N*_0_) vs. Λ(*A, N*). Since the fitness matrix Λ for *N* = 2 depends only on two coefficients, we can represent each of them as points on the plane (*λ*^*j*^(*N*), *λ*^*i*^ (*N*)) and compare their distances. This allows us to quantify the effect of the network *N* by measuring the Euclidean distance between pairs of points in the plane.

Notice that for the homogeneous network *N* _0_ there is an infinite number of matrices *A* yielding the same Λ. We say that two matrices *A*_1_ and *A*_2_ in ℝ^2^ are *homogeneously equivalent* if their corresponding invasion fitness matrices equalize: Λ(*A*_1_, *N*_0_) = Λ(*A*_2_, *N*_0_).

Yet, what we observe in the heterogeneous network is that the resulting Λ depends strongly on the particular choice of *A*, as well as on the variance of the degree distribution. Fig.3 shows an example of the joint impact of both *A* and the standard deviation of the degree distribution *std*(*N*) on the matrix Λ.

We observe that on one hand, for each fixed *A*, there is a strong positive correlation between *std*(*N*) and *d*(*N, N* _0_). Having more variable contact structure in the host population (highly heterogeneous network) increases the distance of the fitness matrix from the homogeneous case, hence impacting more strongly the selection dynamics between strains, and the slope of such sensitivity depends on the particular *A* matrix.

Moreover, for a fixed variance of the network, the details of self vs. non-self interactions. i.e. *A*, impact the Λ of the network not just in a quantitative way (different slope in Figure 3b), but also qualitatively, affecting the directional shift within the 4 quadrants in the plane (Figure 3a).

**Figure 3:**
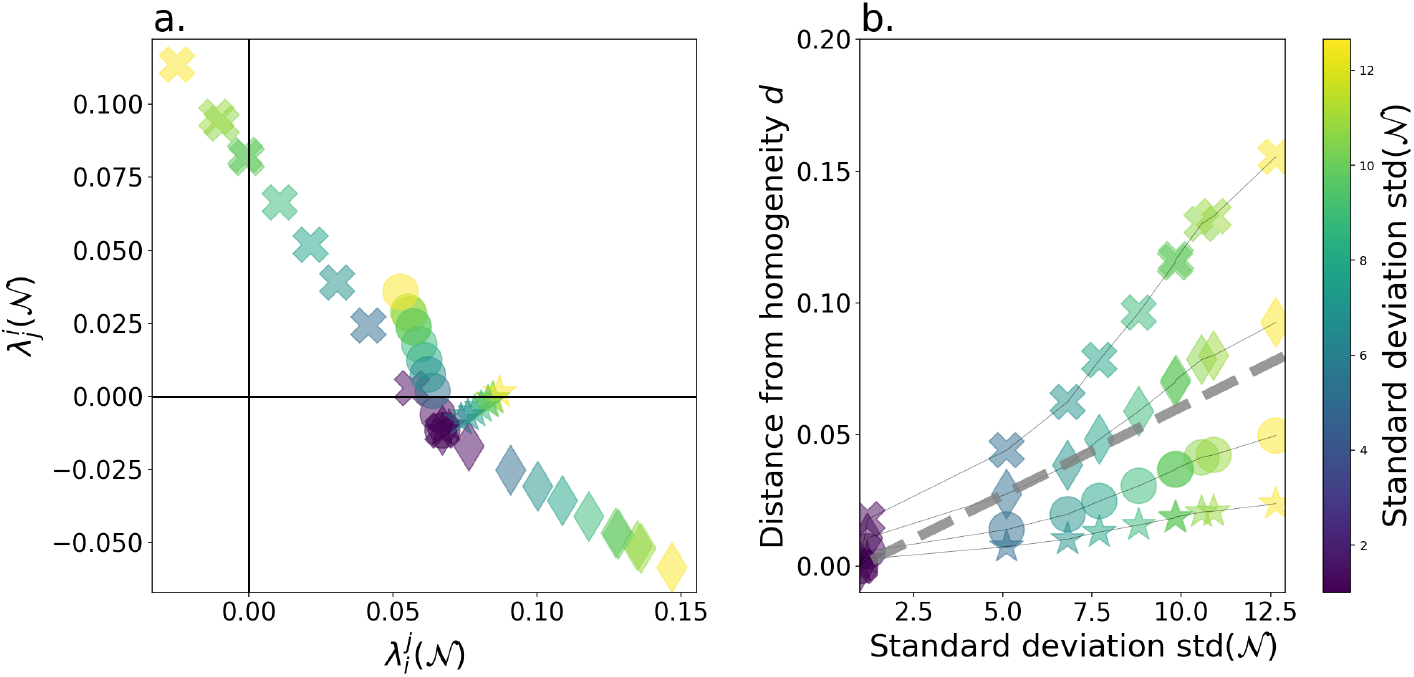
Pairwise invasion fitness changing with network heterogeneity, and absolute distance from homogeneity. On the left sub-figure (panel a.), the pairwise invasion finesses Λ(*A, N*) of two strains *i, j* (*N* = 2) in the plane 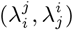 computed from several networks *N* with fixed mean degree ⟨*k*⟩ = 15 and 4 homogeneously equivalent matrices *A* are represented. The colors indicate the variability of the network *std*(N), and the mark indicates the alpha matrices *A*. In black we report the lambda pair Λ(*A, N*_0_) for the homogeneous network (for which *std*(*N*_0_) = 0). Each pair is at a given distance *d*(*N, A*) (as defined in 4.2) from the homogeneous situation. On the right sub-figure (panel b.), we show how the distances for all the networks and all the matrices *A* vary with the standard deviation. The color and the mark are as in the left sub-figure. The *A* matrices are 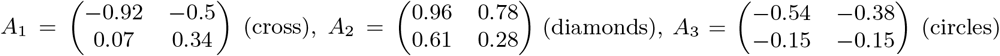 and 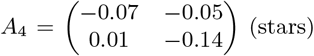. For each choice of the *A* matrix we show 10 networks with different heterogeneity in degree distribution (from blue to yellow). Other parameter values are: *σ* = 1, *β* = 3, *r* = *γ* = 1 so that *R*_0_ = *β/*(*γ* + *r*) ⟨*k*⟩ = 0.15 * 15 = 2.25 in the homogeneous population. The sensitivity of the Λ matrix to network heterogeneity can be viewed in 3 aspects: i) absolute distance from homogeneity, ii) the angle of change in the plane, iii) the qualitative transition between quadrants. In this example, while the underlying *A* matrix is changing, we can see the effect of such architecture of within-/between-strain interactions on Λ −*network* sensitivity in all the three above ways, although in b. only the first aspect is highlighted, the other two remaining hidden. A change in magnitude without *λ*s leaving their starting quadrant implies primarily a change in speed of dynamics (see the diamond case *A*_3_ in a.). A change in magnitude accompanied by a crossing of quadrants implies a change in speed and a change in quality (see the crosses case *A*_1_ in a.). A change in absolute magnitude with network heterogeneity however is a precondition for eventual qualitative shifts in dynamics.

It is well-known that in the *N* = 2 system, the signs of the pairwise invasion fitness coefficients determine the quality of final outcome (see also [22]): e.g. a competitive exclusion scenario favouring strain 1 requires (+,−) sign structure between the two coefficients, while a coexistence equilibrium requires (+, +) coefficients. What we see in Figure 3 is that overall, both the intensity and the direction of the effect of the network on Λ(*N*) depend strongly on the particular matrix *A* within the class of all homogeneously equivalent matrices. How consequential are the resulting network replicator parameters for a *qualitative* shift in strain dynamics, depends on whether the choice of *A* and of the degree distribution heterogeneity are such that the new point 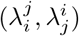shifts quadrant from the original reference point in the homogeneous case. Only in that case, the dynamics would qualitatively shift, e.g. going from an exclusion (+,−) to a coexistence (+, +) scenario; some of such shifts being clearly visible in Figure 3, illustrating the potential of the network to drastically interfere with and overturn selection.

From this first analysis, two interesting metrics naturally arise to quantify the sensitivity of the strain dynamics to the network:

- When comparing *2 networks*: distance between their net lambda’s as coordinates on a plane (*d*)

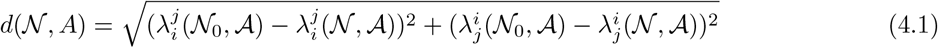
- When comparing *several networks with same mean degree but different variance*: the slope of the linear regression of corresponding lambda distances vs. std of degree distribution:

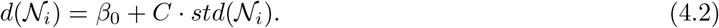

#### The role of mean-field epidemiological parameters and the network

Although hard to see explicitly in the formulas (§2.2.2), the actual network effect on Λ depends strongly on the macroepidemiological parameters: transmission intensity described by the basic reproduction number of the homogeneous model with same mean degree 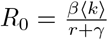 and mean ratio of single to coinfection *σ*, which affect the global and class-specific mean-field epidemiological prevalence of single and co-infection. When we fix the asymmetry in trait structure between strains (*A*), it is interesting to study how these global parameters (*R*_0_, and *σ* ) together with network variance affect the emergent Λ between strains over the network.

In Fig. 4, we show the effect of these two macrolevel parameters for a large number of random networks with the same mean degree ⟨*k*⟩, generated randomly, and pairs of (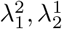) (chosen to explore the four quadrants of the 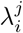 plane, hence forming a circle in the plots). For all these pairs and all the values of *R*_0_ and *σ*, we generate a set of six homogeneously equivalent *A* matrices, and compute the corresponding invasion fitness coefficients over the entire network.

**Figure 4:**
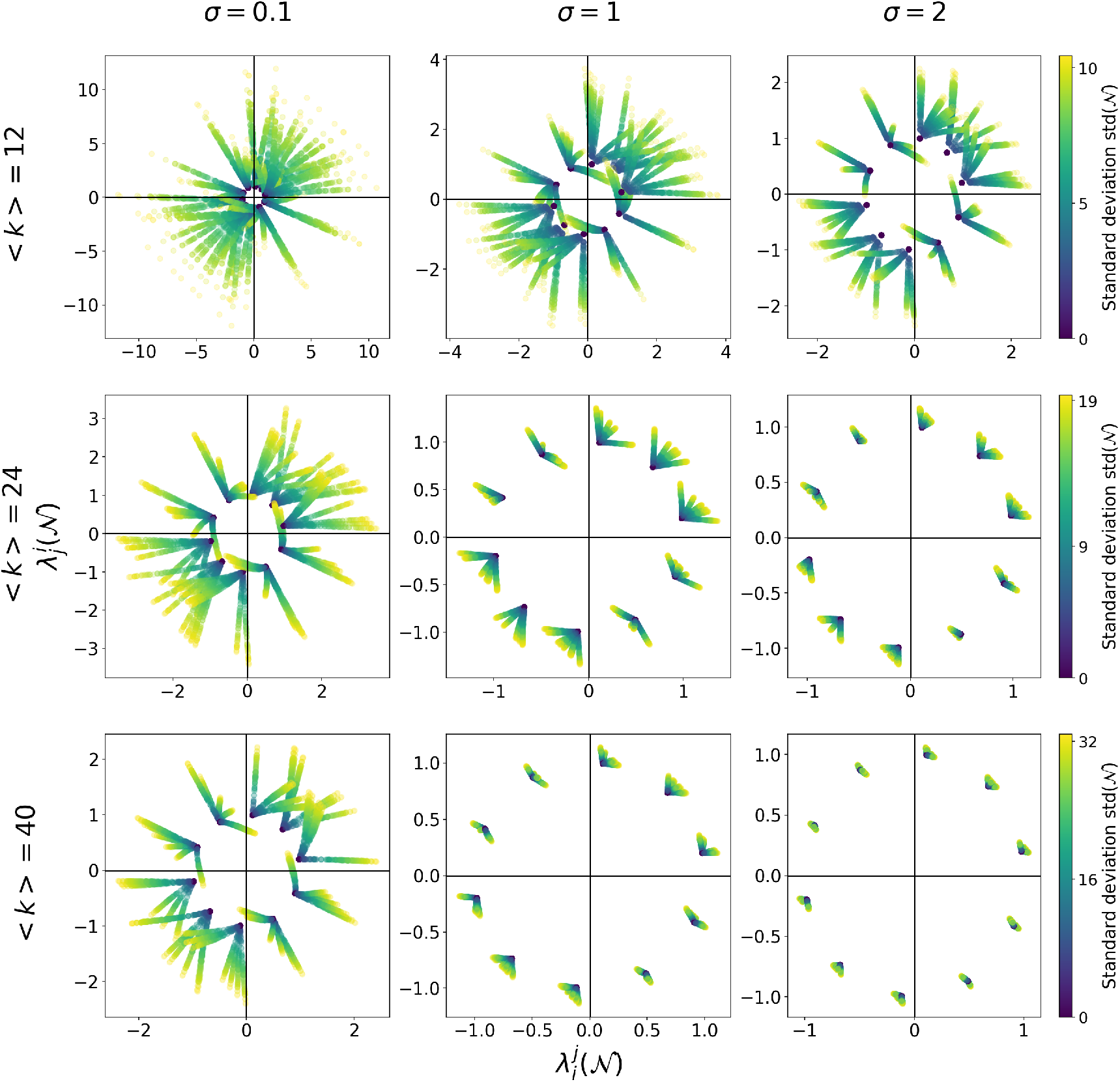
How mean transmission intensity and mean susceptibility to coinfection affect sensitivity of 2-strain dynamics to host contact network structure. We show here the effect of the network’s heterogeneity on Λ(*N, A*) for three different values of transmission intensity, as captured by basic reproduction number *R*_0_ of the homogeneous model (varying ⟨*k*⟩, with *β/*(*γ* + *r*) fixed to the value of 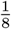 ) and three different values of mean susceptibility to co-colonization *σ*, ranging from overall competition (*σ* < 1) to overall facilitation (*σ* > 1) among strains in co-colonization. All the networks are generated randomly and have the same ⟨*k*⟩, thus the same *R*_0_, row-wise: first row ⟨*k*⟩ = 12 and *R*_0_ = 1.5, second row ⟨*k*⟩ = 24 and *R*_0_ = 3, third row ⟨*k*⟩ = 30 and *R*_0_ = 5. The colors represent the standard deviation of the network’s degree connectivity: in black the homogeneous network, in green-to-yellow colormap the networks with an increasing heterogeneity. For a summary of network effects on Λ in terms of sensitivity measures see Fig.S2.

Our results show that for a fixed value of transmission intensity (e.g. *R*_0_ in the row panels in Fig.4), higher values of co-colonization susceptibility *σ* have the effect of moderating the impact of the heterogeneity of the network. In other words, the more singly-infected hosts can be co-infected by any strain, the less relevant becomes their contact connectivity structure for the strain dynamics over the network, compared to the homogeneous mixing case, indicating that global facilitation in co-infection among strains is less consequential for overall selection on a network than global competition.

Similarly, for fixed values of overall co-colonization susceptibility *σ* (the column panels in Fig. 4), higher values of *R*_0_ tend to reduce the impact of network heterogeneity. In contrast, the closer the basic reproduction number is to the critical threshold value of 1, the more relevant the heterogeneity of the contact network becomes, and so we observe longer radii spreading from the homogeneous mixing reference *λ* origins.

This phenomenon is not affected by the choice of network degree distributions. Varying network structures (Fig. S1) we again observe that host contact network heterogeneity has more impact on Λ for low *σ* and *R*_0_ close to 1, and becomes nearly negligible when at least one of these two macroparameters is large.

It is important to remark that also that in those cases, when the heterogeneity is relevant, i.e. for more competitive co-colonization interactions *σ* closer to 0 or very low prevalence of infection *R*_0_ ≈1, the network effect on Λ, is not only visible in the longer distances from the origin with increasing network heterogeneity, but it can even induce a critical qualitative shift in the results of the replicator dynamics between 2 strains, compared to the homogeneous mixing scenario. Such shift corresponds to network Λ’s crossing the original quadrant of the Λ’s in the homogeneous case.

As clearly visible from the graph, these qualitative shifts come with constraints. Indeed they are more likely to occur: from an exclusion regime to a coexistence or bistable regime, but not from one exclusion to the other exclusion and not from stable coexistence to bistability.

This is visible in the apparent *butterflies* in the 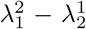 plane in Figure 4 always pointing in opposite directions with respect to the left-right diagonal.

From equation 2.12 and the definition of the pairwise fitness 2.13, it is possible to gain insight on this effect of the network on Λ. In fact, the only effect that the weight Ξ_5_ can have on 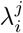 is to enlarge the cloud of points in the plane, while *µ* (the global ratio of single to coinfection) dictates the direction, and will only affect the antisymmetric part. In other words, *µ* is a global quantity, and as we showed in [21], it appears to be always lower in the contact network than in its homogeneous counterpart *µ*(*N* ) *< µ*(*N*_0_). Increasing the network heterogeneity increases the symmetric part of Λ and decreases its antisymmetric part. This means that, irrespectively of the network or its heterogeneity, no mechanism allows the strain dynamics to cross from coexistence to bistability or vice versa, or from one exclusion to the opposite. This is to be expected from this trait-based model version of Λ (Eq.2.16), where the only strain variation is in co-colonization vulnerabilities.

#### Interplay between different parameters in concise sensitivity measures

How can we summarize the information in graphs such as those in Figure 4? In order to quantify the combined effect of global parameters and network heterogeneity, we focus on the sensitivity to the contact network variance as expressed by the *C* measure (see Fig.S2). We observe that it decreases with mean susceptibility to coinfection *σ*, for all values of transmission intensity. However, the effect is notably bigger for smaller *R*_0_, consistent with length of radii in Figure 4. This means that under fixed homogeneously-mixing multi-strain systems, the more cooperation there is on average in co-infection within and between strains, the smaller will be the effect of host contact network on the strain dynamics, and viceversa: the more competition there is on average in co-infection, the more relevant will become the heterogeneity in host contact networks for ultimate strain dynamics. When varying the topology of the network, we see the pattern remains similar, but that this coefficient is higher in the random topology than in the three-class network case (Fig.S2 b. vs. a).

### 4.2 Impact of network heterogeneity - the case of *N >* 2 strains

Until now we have considered the simple, yet nontrivial case of just 2 strains co-circulating on a host contact network. Next we explore how the host contact network degree distribution impacts circulation and selection dynamics between a higher number of strains.

#### Impact of the network heterogeneity on the *N* -strain fitness matrix

In this section, we consider the matrix Λ for *N* = 10 interacting strains and examine the impact of network heterogeneity. Again, we fix all epidemiological parameters and the alpha matrix *A* ∈ *ℳ* _*N*_ while we change the network.

In Fig. 5, we show how the network heterogeneity impacts qualitatively and quantitatively the invasion fitness values and the dynamics of the strain frequencies as represented by the replicator system. The three columns in the figure correspond to different network structures. In the upper row, we show the host contact degree distribution. The first case corresponds to the strictly homogeneous degree network (for which *std*(*N*) = 0), the second corresponds to a heterogeneous network with 3 degree classes, symmetric around the mean, and the last represents an arbitrary random network. They all have the same average degree (⟨*k*⟩ = 20). The second row shows the pairwise fitnesses for all pairs of strains (different points) for each network structure.

**Figure 5:**
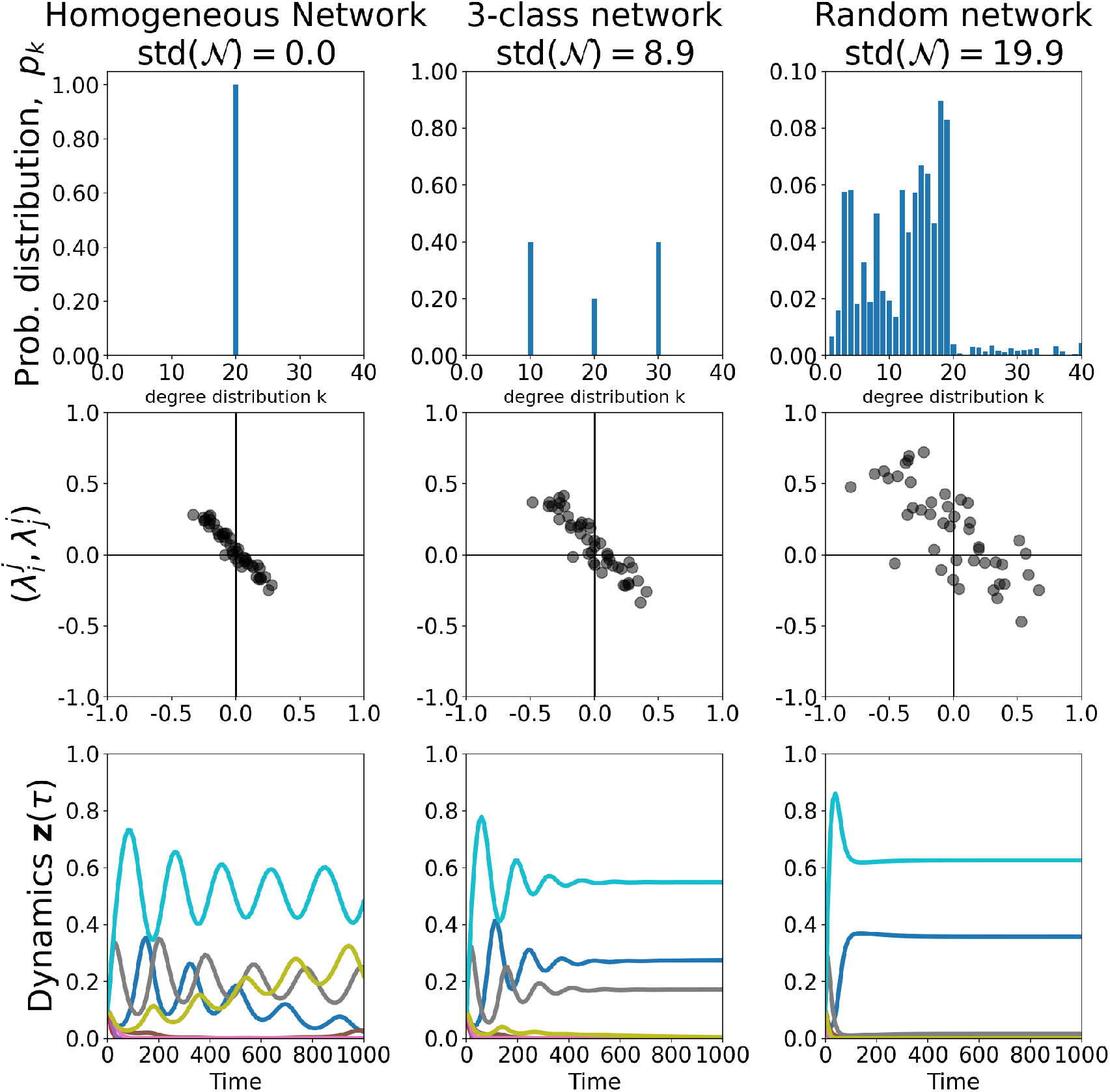
Network heterogeneity impact on invasion fitness matrix for *N* strains. We show three networks with average degree ⟨*k*⟩ = 20: *N*_0_, *N*_1_, *N*_2_ respectively the homogeneous network (where all nodes have degree ⟨*k*⟩ = 20 and hence *std*(N) = 0), a network with only three degree classes (*k* ∈ {10, 20, 30} with probabilities {0.4, 0.2, 0.4} and *std*(N) = 8.9), and last a network with 97 degree classes (and *std*(N) = 19.9). Here *β* = 0.15, *r* = 0.5, *γ* = 0.5, ⟨*k*⟩ = 20 and *σ* = 0.3 so, for the homogeneous network, 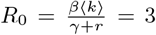. We plot the degree distribution, the pairwise fitness values, and the replicator dynamics respectively in the first, second, and third row panels. In the dynamics all strains start from equi-abundance.

For each network *N*, we notice that typically the pairwise fitness values 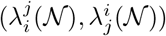 spread over the four quadrants, and the effect is stronger for larger *std*(*N*). More heterogeneous networks increase the space covered by pairwise fitness coefficients, increasing thus the diversity of invasion network edges (+ +: coexistence, - +: competitive exclusion, - - :, bistability, as described in [11, 22]).

This is expected to make the dynamics less complex pushing the possible outcomes of the replicator dynamics towards the more stable regimes [11]. We plot corresponding dynamics in the last row. Notice how in the homogeneous network (bottom-left plot) we observe strain oscillations for a long time without reaching any stable equilibrium. Increasing the heterogeneity (bottom-central plot) shortens the time for the stabilization of the strain frequencies. Increasing further the heterogeneity leads to even faster stabilization and also (in this particular case) a reduced number of persistent strains.

These simulations seem to suggest that the existence of host contact heterogeneity can stabilize otherwise complex multi-strain dynamics in SIS endemic scenarios with strain interactions.

#### Impact across a broad range of epidemiological settings, pathogen diversities, and host network structures

As seen in the previous section, host contact heterogeneity can affect drastically the replicator dynamics. How generalizable are these results? A distinctive feature of the replicator equation is that the system is linear in the interaction matrix, given in our case by the strain pairwise invasion fitness matrix Λ. As a consequence, if two interaction matrices differ only by a positive multiplicative constant, Λ_1_ = *s* Λ_2_, then the *quality* of the dynamics remains exactly the same, while the *speed* of the dynamics is simply rescaled by the factor *s*. Motivated by this observation, we decompose the system into two components: a measure of the magnitude of the interaction matrix, denoted *Speed*(*N*), and a renormalised matrix Λ_norm_(*N* ) that captures the intrinsic structure of the dynamics independently of its overall speed.

Note that, since deviations from neutrality are assumed to occur only through susceptibility to co-colonization, the factor Ξ_5_(*N*) affects only the overall timescale of the dynamics and disappears after renormalization. Consequently, network structure can influence the qualitative dynamics only through the parameter *µ*(*N* ) (see Section S1.1).

In this section, we examine how the contact network influences two key quantities: (i) the speed of the dynamics and (ii) the *dynamical distance* of multi-strain dynamics on the network from the corresponding dynamics in the homogeneous population. The latter is assessed by studying (iii) the distance between the normalised interaction matrix and its homogeneous counterpart.

To cover as many parameter regimes as possible, we compute the replicator dynamics over 1000 networks with various types of topology (3-class symmetric, fully random, uniform, and binomial) with different strain number *N*, different randomly-generated strain interaction matrices *A*, mean contact number ⟨*k*⟩ and mean coinfection vulnerability factor *σ* . All the primary parameters are shared between the homogeneous and heterogeneous case, except the fact that the network involves a distribution over contacts given by the random *P* (*k*) and corresponding *std*(N) each time.

We study how the heterogeneity of the network affects the matrix Λ(*N*) compared to the homogeneous situation Λ(*N*_0_), from which we can derive information on the speed of the dynamics after suitable normalization (see Table 2). In Fig.6a we show that the speed of the dynamics in the case of heterogeneous network relative to the homogeneous topology increases with the standard deviation of the network (normalized by the mean degree). For the linear relation, we derive 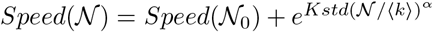, highlighting that increasing the coefficient of variation in host contact networks speeds up multi-strain dynamics.

**Table 2:**
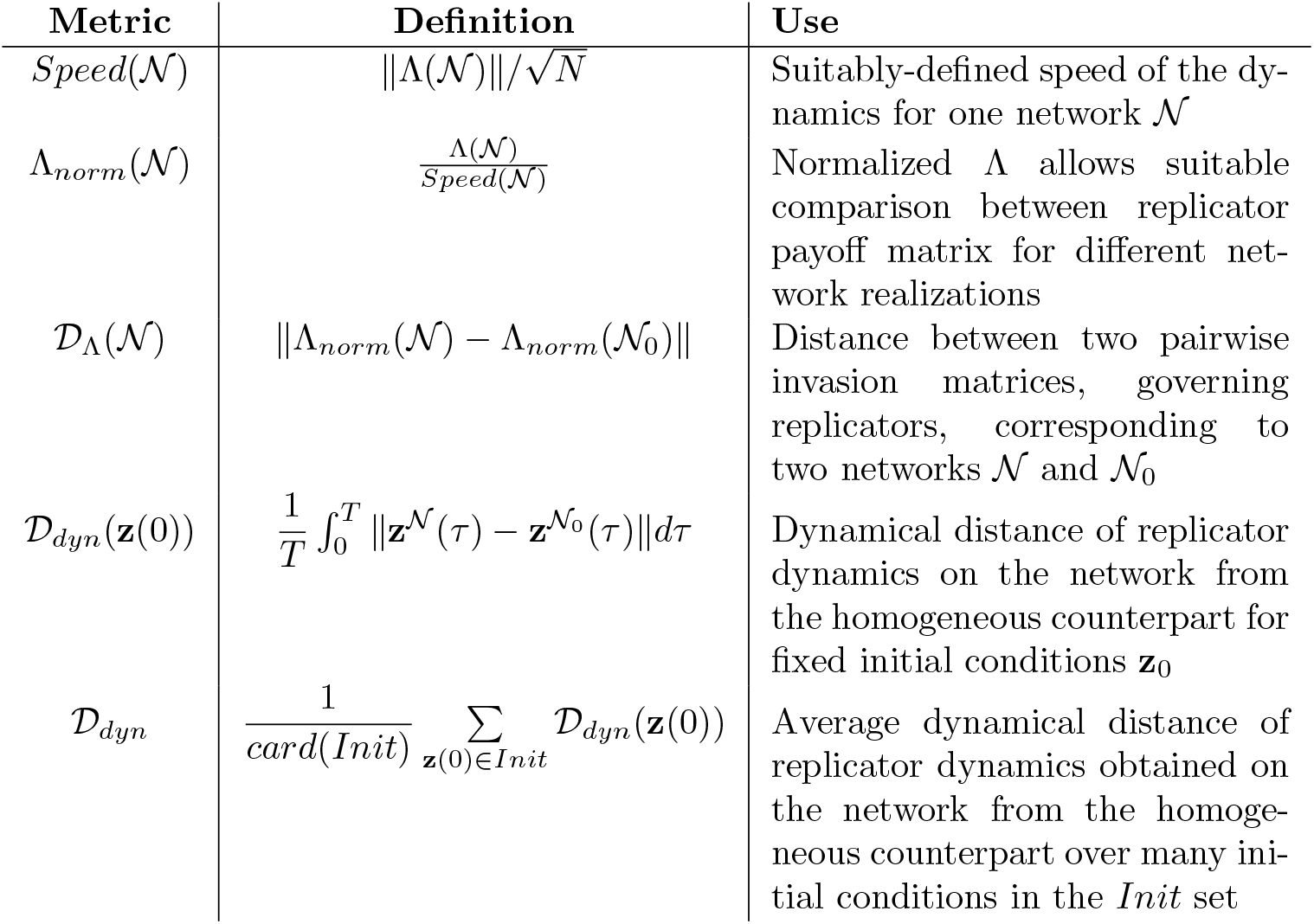
Metrics used to quantify the effect of the host contact network on multi-strain dynamics.

To better evaluate the impact of heterogeneity we study also explicitly the quality of the dynamics via system simulations. We proceed as follows. In each of the 1000 systems, for several initial data, we compare the actual trajectories 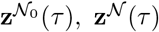 between the homogeneous network and the heterogeneous one, using dynamical distance metrics, then we take the mean distance starting from several initial conditions **z**(0) ∈ *Init* that are identical for the two networks. This leads to what we call *dynamical distance from homogeneity D* _*dyn*_ (see Table2). Figure 6b indeed shows that this dynamical distance between the two replicators increases with network coefficient of variation, hence confirming that the heterogeneity of the network does quantitatively affect the multi-strain dynamics.

**Figure 6:**
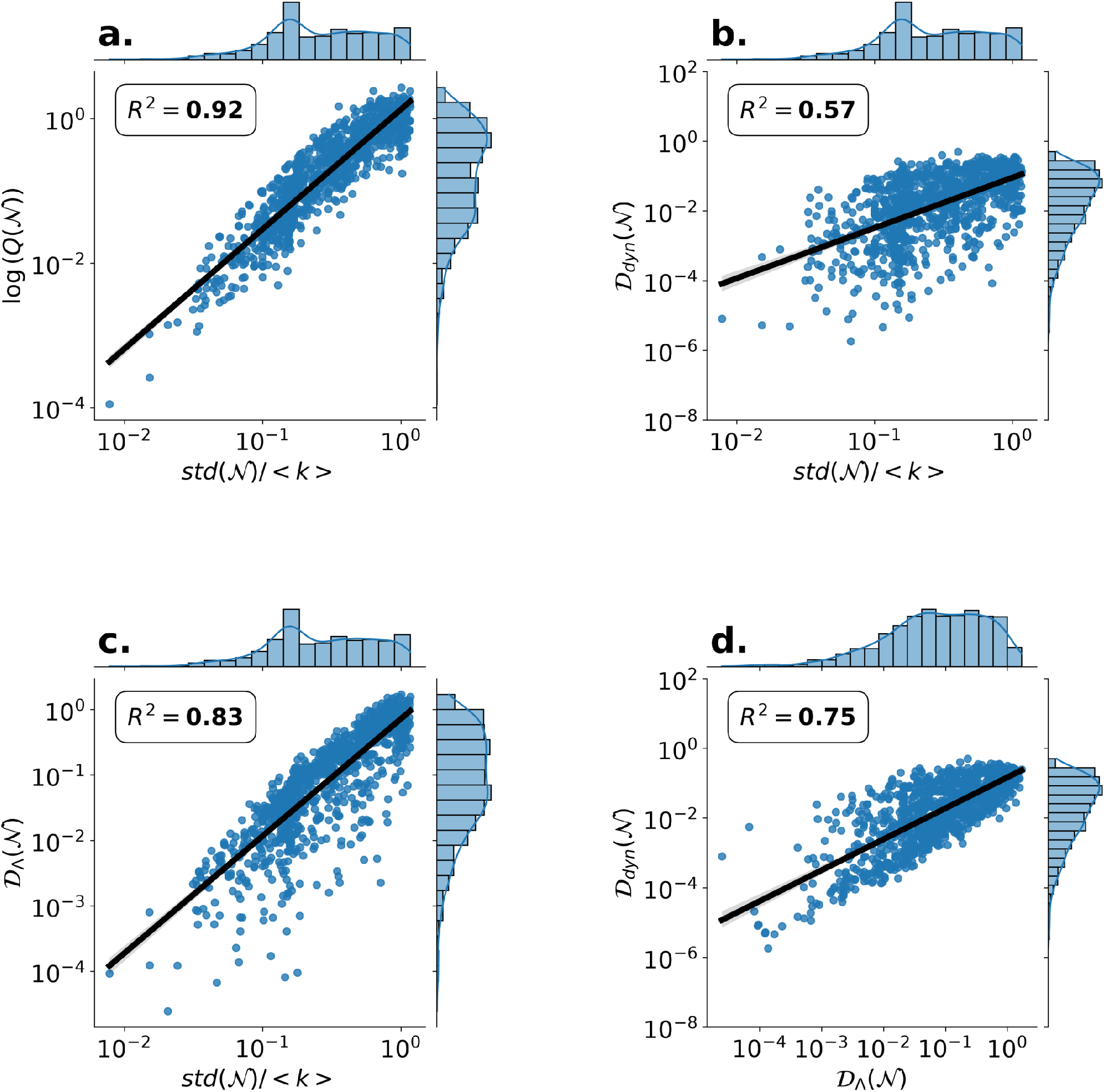
Network heterogeneity impact on the quality of the multi-strain replicator dynamics. We generate 1000 replicator systems with different heterogeneous networks, different numbers of strains *N*, different biological parameters, and different alpha matrices *A*. Each time, we compare metrics relative to the corresponding homogeneous network case with the same mean degree. All the axes are presented in log-scale. The quality of linear fits is represented through the *R*^2^ coefficient for each metric. Parameter values satisfy: *σ* ∈ (0, 2); *β/*(*r* + *γ*) = 0.11*/*(1 + 1) = 0.055 ; *< k >*∈ {20…30}, and *N* ∈ {2, …, 9} and *T* = 1000. **a**. We represent the log of the average speed of the dynamics for a network *N* relatively to the homogeneous network 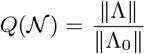 as a function of the coefficient of variation (standard deviation of the network degree distribution normalized by the mean degree) *std*(*N*)*/* ⟨*k*⟩. **b**. We represent how the dynamical distance from homogeneity D_*dyn*_(*N*) (computed over *N* + 1 different initial conditions for each system, one uniform and *N* close to each corner *z*_*i*_ = 1) varies with *std*(*N /* ⟨*k*⟩). **c**. We represent how *D*_Λ_(*N*) depends on *std*(*N*)*/* ⟨*k*⟩ **d**. We show that a significant part of the variation in dynamical similarity D_*dyn*_(*N*) can be well-captured by replicator Λ-matrix similarity, *D*_Λ_(*N*). This suggests that the computation of corresponding matrix distances D_Λ_(*N*) - which is easy and quick - is a good first approximation to quantify the effect of network heterogeneity on multi-strain dynamics and compare different networks. For a stratified version of this figure by the epidemiological parameter *σ* denoting vulnerability to coinfection, see Figure S3.

Unsurprisingly, as expected, such dynamical distance is a direct result of the Λ matrix distance. Indeed, we observe that the distance *D*_Λ_(*N*) defined in Table2, also increases with the standard deviation of the contact network (Figure 6c). We argue that such distance explains well the difference between the qualitative dynamics in a network *N* and the corresponding homogeneous network *N*_0_ (Figure 6b). To support this statement, in Figure 6d, plotting them against each other, we show that the dynamical distance of the multi-strain system increases with the distance between the normalized strain invasion fitness matrices of the heterogeneous and homogeneous networks. The results hold generally and robustly for all the network structures studied in this work. We conclude that the heterogeneity of the networks explains well the differences in the strain dynamics by affecting the overall speed, as well as the quality of the replicator through the matrix Λ_*norm*_.

When stratified by the specific value of the interaction coefficient in co-colonization among strains, *σ*, and tested within a formal regression (OLS) framework, the simulations in Figure 6 reveal clear patterns highlighting the effect of the epidemiological parameter *σ* (see Figure S3).

#### Impact on the number of coexisting strains

From the previous results, it is clear that heterogeneity in the contact degree distribution strongly affects the speed of multi-strain propagation on host contact networks and the quality of their dynamics. And it does this via changing the payoff matrix of the corresponding replicator system, namely the pairwise invasion fitness Λ matrix. Although dissecting all the effects on multistrain dynamics would require more in-depth analysis of specific scenarios and explicit accounting of stability-complexity relationships, here we will focus only on one additional aspect: the number of coexisting strains. The number of coexisting strains is a very basic measure of diversity, which does not depend on the precise values of the *z*_*i*_. This is very convenient in our situation, as it allows us to deal with potentially complex asymptotic dynamics such as cycles or strange attractors. Hence the principle is to take *T*_max_ large enough to reach the attractor and avoid transient dynamics, and then to compute the number of indices *i* such that *z*_*i*_(*T*_max_) ≠ 0. In practice, however, several technical issues arise, detailed below.

First, if *z*_*i*_(0) *>* 0, then *z*_*i*_(*t*) *>* 0 for all *t*. We therefore need to relax the measure by counting the number of strains above an arbitrary fixed small threshold t = 0.1*/N* . Second, the condition “*T*_max_ large enough” is not precise: the required time strongly depends on the particular system. We therefore first renormalise Λ(N) by its speed *Speed*(*N*), which yields the same average time unit for each system, and then we take the arbitrary value *T*_max_ = 5000, which seems to be sufficiently large in most situations. Third, the measure is consistent only if the persisting strains on the attractor stay far from the boundary *z*_*i*_ = 0. This holds when the persisting strains are permanent - that is, they belong to a compact subset of {*z*_*i*_ *>* 0} - but not always. For instance, if the attractor is a union of heteroclinic cycles, some strains may remain at very small values for a long time before returning to the system. Our measure does not capture diversity well in such cases. Lastly, the measure depends on a single trajectory starting from the particular initial condition **z**(0) = (1*/N*, …, 1*/N* ). It therefore only captures the number of persisting strains for the attractor reached from this initial condition, and misses strains that may persist when starting from other values (see [26] for a deeper discussion of this issue). This must be kept in mind when the system exhibits multiple attractors; which may be the case here for small *µ*, see for instance [11].

Using such rationale, in Figure 7 we show one example of the effect of network heterogeneity on the number of interacting strains persisting in a system. We generate 2500 random networks in each of 4 categories of degree standard deviation (*std*(*N*)), and simulate multistrain dynamics on these networks up to a fixed, sufficiently large, time *T* = 5000, starting from strain equi-abundance (*N* = 20). We observe for these parameters (mean degree ⟨*k*⟩ = 12 and coinfection vulnerability factor *σ* = 0.5) that the higher the heterogeneity in the host contact network, the smaller the number of coexisting strains, all else equal. The highest mean number of persisting strains is observed in the homogeneous network case where all hosts have the same degree. Such reduction effect of host contact network heterogeneity on strain diversity is similar to what has been found in previous multi-strain models [33]. Yet, when varying parameter combinations (Figs.S4 and S5), we notice a more complex bigger picture: this effect of contact network on the number of persisting strains can be stronger, weaker or almost null, depending on the global epidemiological parameters driving transmission intensity and coinfection, such as ⟨*k*⟩ and *σ* . In particular, somewhat greater diversity reductions are observed for lower values of *σ* and *k*, where the epidemiological system tends towards regimes of more competition among strains and lower transmission intensities. Remarkably, these are precisely those regimes expected to exhibit more complex multi-strain dynamics in the corresponding homogeneous populations [11] to start with. It seems that in such cases, by lowering *µ* (Fig. 2 e-g), similar to what we have described earlier in [11], the network-induced heterogeneities shift the dynamics towards regimes of more niche effects, stability, and fewer strains coexisting.

**Figure 7:**
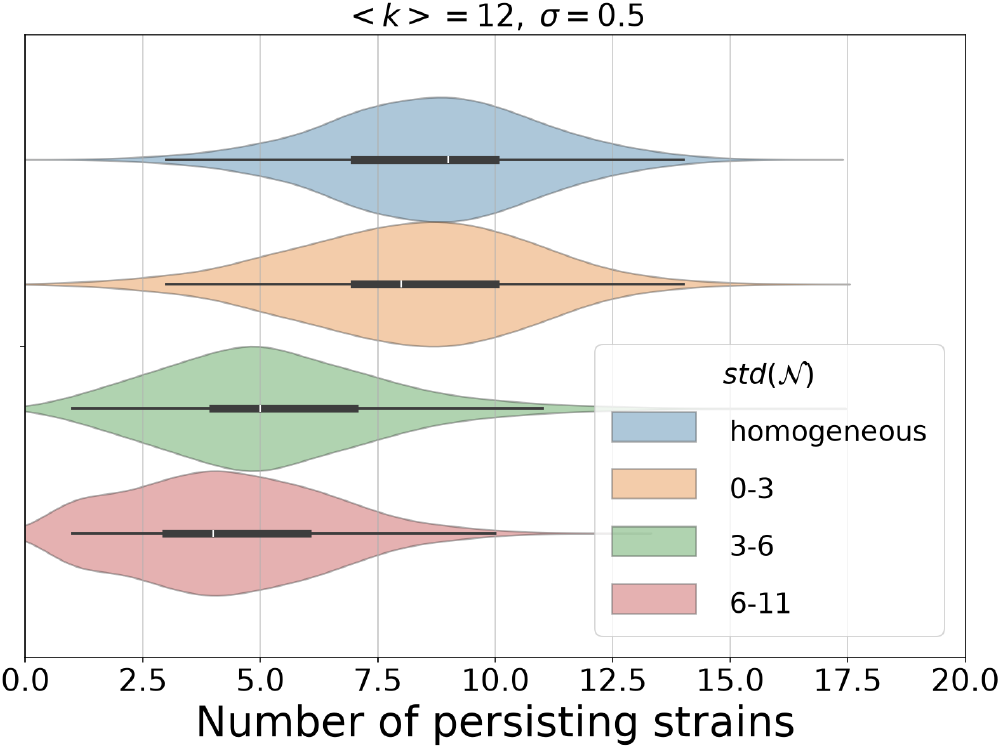
Number of persisting strains in SIS dynamics with co-infection depends on host contact network heterogeneity. We show violin plots of the distributions of the number of persisting strains over 10.000 replicator systems (§2.2.2. Eq. 2.12 and 2.16), for *N* = 20 strains, grouped by range of *std*(*N*). These were obtained by simulations on randomly generated networks and under random strain deviation from neutrality (*α*_*ij*_ coefficients). For each system, we compute the dynamics starting from equi-abundance 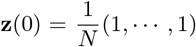 and ending at *T* _*max*_ = 5000. The final output is computed from **z**(*T*_*max*_) = (*z*_1_,*·*··, *z*_20_). The number of persisting strains is defined by the number of indices *i* for which *z*_*i*_ *>* 0.1*/N* . Parameter values: *σ* = 0.5, ⟨*k*⟩ = 12 and 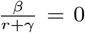 thus *R*_0_ = 1.5. We group the 10 000 systems in 4 categories of equal size (2500 each) according to the value of *std*(N). Group 1 (blue): *std*(N) = 0 (Hence, there is only one class with *k* = 15; this is the homogeneous system). Group 2 (orange): 0 *< std*(*N*) ≤ 7. Group 3 (green): 7 *< std*(*N*) ≤ 15. Group 4 (red): 15 *< std*(*N*) ≤ 25. For sensitivity of these results to different parameter values ⟨*k*⟩ and *σ*, see Figs.S4 and S5 in the Supplementary Section S2.

#### Coinfection vulnerability *σ* and overall transmission intensity *R*_0_ as mediators of network effects

To understand more deeply the above effects, we must recall that both *σ*, 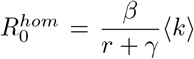 and std(*N* )*/*⟨*k*⟩ influence *µ* (Figure 2) and thereby the invasion fitness coefficients of the replicator through

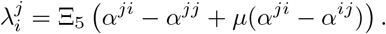

Wherein we have set the common factor Ξ_5_ = 1 for simplicity without affecting the *qualitative* description. Since *µ* decreases with both std(*N* )*/*⟨*k*⟩ and *σ*, the magnitude of 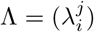 decreases as *σ, R*_0_ or std(N) increase. For large *σ*, the baseline value *µ*_0_ is already small, so std(N) can only weakly affect *µ*, and thus the global amplitude 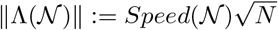 varies little. We highlight two effects as follows.

i. *Effect 1*. When *σ* (and *R*_0_) increase, *µ* decreases uniformly with respect to std(*N* )*/* ⟨*k* ⟩, making the impact of heterogeneity on the magnitude of Λ(*N*) small. The mean dynamical speed simply reflects this amplitude *Speed*(*N*). By [11],this explains why network heterogeneity tends to favour faster dynamics with many stable equilibria and low biodiversity (Figure S4 S5), and why this effect becomes negligible for large *σ* and *R*_0_. Consistently with this, for fixed *σ* we find that higher transmission reduces the influence of network heterogeneity on selection, whereas lower transmission (especially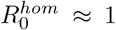) increases it. Heterogeneity is therefore most critical near threshold conditions, where small structural differences in the contact network can tip the balance between competing strains, while for large *σ* or *R*_0_ the multi-strain dynamics approximate those of homogeneous systems. However, turning to the *quality* of the dynamics, only the renormalised matrix Λ_*norm*_(*N*) = Λ(*N*)*/Speed*(*N*) matters. The distance *D*_Λ_(*N*)between Λ_norm_(*N*) and its homogeneous counterpart Λ_norm_(*N*_0_) grows with both std(*N*)*/* ⟨*k* ⟩and *σ* (Figure S3), showing that heterogeneity has a stronger dynamical effect when *σ* is large, opposite to Effect 1. The mechanism is structural. Network heterogeneity acts only through the antisymmetric term *α*^*ji*^ −*α*^*ij*^, so its influence is always directed along the antisymmetric axis. If the homogeneous system is already nearly antisymmetric, heterogeneity merely reinforces this and renormalisation largely suppresses the effect. In contrast, if the pairs 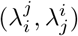 are predominantly symmetric, then std(N) pushes the system toward the antisymmetric direction, producing a substantial change in the distribution of the 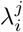, even after renormalisation (Figure S6). This situation occurs precisely when *σ* is large: *µ* is then small, and the homogeneous interaction matrix exhibits a high symmetry index. In this case, the system is not yet aligned with the antisymmetric direction, so heterogeneity has more leverage to rotate the interaction pairs toward that direction. As a result, heterogeneity more strongly modifies the renormalised matrix Λ_norm_(*N*) and thus the dynamics.
ii. *Effect 2*. When *σ* is large, *µ* is small and the homogeneous matrix is more symmetric. Consequently, network heterogeneity is more likely to rotate the interaction pairs toward the antisymmetric direction and therefore to modify the renormalised matrix Λ_norm_(*N*). This explains why the dynamical variation induced by heterogeneity increases with *σ* . In summary, increasing *σ* or *R*_0_ produces two opposite effects:
  i. a reduction in the *magnitude* of variations (Effect 1), and
  ii. an increase in the *susceptibility* of the system to rotation toward the antisymmetric direction (Effect 2), because the homogeneous pairs are more symmetric.

Tests on multiple networks consistently show that the interplay between the intensity of heterogeneity and the epidemiological parameters *σ* and 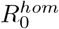 is robust across different network structures (Supplementary Figs. S1–S6), underscoring that the observed effects are not topology-specific but a general property of multi-type dynamical processes in structured populations [21].

Overall, our results emphasize that network heterogeneity can now be explicitly embedded and studied in models of co-infection and multi-strain dynamics, with potentially strong effects, especially in competitive systems, and transmission threshold regimes. The architecture of strain traits, and in particular, the specific form of within- and between-strain cooperation and competition, interplay with network structure to determine the selection outcomes, giving rise to a complex context-dependent picture. Although a simple rule of thumb may remain elusive, the proposed reduction of multi-strain SIS dynamics with coinfection to a global replicator on a host contact network offers a much simpler way to investigate specific effects. These can be examined case by case or through appropriate stratification, primarily by comparing pairs of replicator systems. These insights have broad implications for understanding the persistence, coexistence, or extinction of pathogen strains in realistic, heterogeneous populations, and potentially for designing interventions that exploit network structure to control endemic infectious diseases.

## 5 Discussion

We have presented a framework for modeling and investigation of coinfecting multi-strain SIS dynamics on host contact networks. To our knowledge, this study is the first to propose a direct formal mapping between replicator dynamics and multi-strain epidemiology on networks. By focusing on the emergent global replicator dynamics on the network [21], and its constituent net parameters, our results demonstrate that network heterogeneity plays a pivotal role in shaping the selection dynamics between strains in endemic systems with co-infection and interactions. A major simplifying assumption was strain similarity, hence our replicator model reduction results apply in quasi-neutral multi-strain systems, yet, as shown in our previous work, the slow-fast dynamics approximation remains accurate also in regimes with less strict similarity, i.e. when *ϵ* is not too small [18], and thus may reach wider applicability.

### The impact of network heterogeneity

A novelty of our framework is the explicit emergence of the particular payoff matrix, in our case invasion fitness matrix between strains (Λ) in the global replicator governing strain selection on the network (§2.2.2). This allows to compare directly the resulting matrix with the corresponding benchmark in the homogeneous unstructured population. Previous theoretical work has also studied evolutionary games on regular graphs, showing that moving a game from a well-mixed population onto a regular graph simply results in a transformation of the payoff matrix of the replicator equation [28].

In our case, this transformation of the payoff matrix in the replicator equation arises in a nontrivial way from network heterogeneity and strain heterogeneity, modulated by global mean-field epidemiological quantities. Although the general replicator allows for variation along 5 dimensions between strains, we focused on the simplest scenario of variation only in co-infection vulnerabilities [22, 11]. What emerged from our results is that such replicator dynamics is sensitive not only to the structure of the network itself, i.e. degree variance, but also and crucially to i) the specific underlying architecture of the self/non-self strain interaction traits (standardized matrix *A*) and ii) global mean-field epidemiological quantities such as global transmission intensity, or mean-field facilitation coefficient between strains in co-infection.

We show that both the intensity and direction of the network’s impact on Λ(*N*) depend strongly on the particular choice of *A*, even when *A* belongs to the class of homogeneously equivalent matrices. This indicates that interaction matrices which yield identical dynamics in a well-mixed or homogeneous setting can diverge dramatically when embedded in heterogeneous networks. As such, depending on underlying interaction structure, increasing network heterogeneity can induce fundamentally different selection outcomes between strains, including shifts from epidemiological strain exclusion to co-existence or bistability.

### Influence of global epidemiological parameters

In our model all three factors: global coinfection vulnerability *σ*, - an index of general cooperativity among strains, transmission intensity 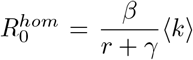, and network heterogeneity, quantified by std(*N*)*/*⟨*k*⟩, influence *µ* and consequently the strain invasion fitness coefficients in the replicator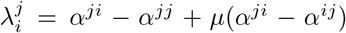. Increasing *σ* or 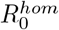 decreases *µ*, reducing the magnitude of the interaction matrix Λ(*N* ) and weakening the influence of heterogeneity on the mean dynamical speed. This is consistent with the tendency of heterogeneous networks to promote slower dynamics, multiple stable equilibria, and lower biodiversity [11], an effect that becomes negligible for large *σ* and 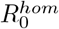 (Figure S2, S5). Conversely, heterogeneity is most influential near the epidemic threshold, where small structural differences can alter strain selection. However, we showed that considering the renormalised invasion fitness matrix Λ_norm_(*N* ) = Λ(*N* )*/Speed*(*N*), the dynamical effect of heterogeneity increases with both std(*N*)*/* ⟨*k*⟩ and *σ* (Figure S3). This apparent contrast arises because heterogeneity acts through the antisymmetric component *α*^*ji*^ −*α*^*ij*^, promoting a rotation of interaction pairs toward the antisymmetric direction. When *σ* is large (promoting more coinfection), *µ* is small and the homogeneous population invasion fitness matrix is more symmetric, allowing heterogeneity to induce a stronger modification of the renormalised matrix and thus of the dynamics (Figure S6). Therefore, increasing *σ* or 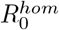 simultaneously reduces the magnitude of dynamical variations while potentially increasing the system’s susceptibility to network structural heterogeneity. These effects are robust across different network structures (Supplementary Figs. S1–S6), suggesting that they reflect a general property of multi-type dynamical processes in structured populations rather than a topology-specific phenomenon [21].

### Metrics for quantifying network effect

Crucially, the network effects on multi-strain dynamics in our SIS framework with coinfection are not merely quantitative, but they can qualitatively alter the ultimate dynamics. We observed that increasing contact network heterogeneity can cause the pairwise fitness between two strains to cross quadrant boundaries, dramatically changing selection outcomes. For instance, systems that would otherwise exhibit single-strain dominance under homogeneous conditions may allow co-existence or develop bistability under increased heterogeneity. In systems with more than 2 strains, accessible now under our formalism, such pairwise transitions would have strong effects on overall *N* -strain dynamics.

To systematically quantify these effects, we introduced several metrics. For example, for *N* = 2, the sensitivity index *C* measures the regression slope of network lambda distances from the homogeneous lambdas vs. network variance. We showed that *C* consistently decreases with *σ*, with stronger effects at lower *R*_0_.

We also proposed three measures of speed and dynamical distance to compare multi-strain dynamics on heterogeneous networks vs. in homogeneous populations: *D*_Λ_, *D*_dyn_, and *std*(*N*) for multi-strain systems of arbitrary size. Examining their co-variation across many systems and for large strain numbers we obtained results confirming that network heterogeneity impacts quantitatively, but also the qualitative nature of the dynamics. Indeed, the effect on the speed is to typically increase it, accelerating selection processes, whereas the effect on the quality of the dynamics seems to be a stabilizing one that decreases the number of coexisting strains, although generalizations warrant further investigation.

### Links with other models and future outlook

Models for multiple diseases that co-spread in a host contact network have gained much attention in recent years [6, 15, 37], even extending the study to multilayer networks [7, 8, 9, 16, 36]. Although our work resonates with some of these previous studies in general spirit, there are definite differences in model structure, findings and scope. For example, we include same-strain coinfection [22, 1, 38], which other models do not typically include (often focusing on two different diseases [15]). We define global susceptibility to coinfection (*σ* ) as an index of global cooperation among strains on a continuum spectrum from values below 1 to values above 1. In contrast, other works define cooperative contagion as a trait only between two different diseases or strains [6]. We use a strain similarity assumption which allows to simplify the model representation into a fast and slow component, yielding an explicit replicator equation between strains - not available with existing multistrain models on networks. We study the standard deviation (and coefficient of variation) of the host degree distribution as a measure of network heterogeneity and bifurcation parameter for strain selection, while other works focus on other properties of the network such as clustering coefficient [15] and seek critical thresholds for total transmission and overall prevalence (summing over diseases) instead of each specific disease. For example, [6] investigate the SIS case with 2 cooperating diseases, comparing deterministic well-mixed and networks 2d-lattice system and Erdos-Rényi networks. They find and highlight discontinuous transitions in overall prevalence mediated by the magnitude of the cooperative coefficient.

Despite the differences in details of model formulation and exact findings, on a broad level, our work - primarily addressed to describe and compare selection dynamics, - also shows that network properties can drastically affect interacting multi-type disease transmission. Host contact network architecture can tip both overall transmission and the balance of strain selection, with network heterogeneity acting to lower the threshold for overall infectious disease persistence, but also typically tending to stabilize and reduce diversity in multi-strain dynamics (Fig. 7). The effect of diversity reduction in host contact networks has been suggested also in previous multi-strain models [33].

With regards to the effects of cooperation, we find that in multi-type systems displaying higher overall cooperation coefficient in co-infection, final strain dynamics, as captured by the replicator Λ matrix, are less sensitive to host network structure, and epidemiological selection should settle closer to what’s expected in the homogeneous case. Conversely, when coinfection vulnerability is lower among all strains (*σ* < 1), hence increasing overall competitiveness in the system, the host contact degree distribution interferes more strongly with strain selection. The explicit nature of the global replicator equation governing strain dynamics on networks, presented here, should make this aspect, as well as other qualities of strain dynamics (diversity, complexity, stability) amenable to proper and full analytical treatment in the future.

### Limitations and further considerations

Due to the mean-field assumption used here to represent the host contact network, the results should be interpreted as a first-order approximation rather than a comprehensive account of network effects. As mainly done in the literature, the mean-field approach zooms out and loses track of individual connectivity. Full information on individual connectivity would lead to a more precise analysis. In the future, it will be interesting to verify these results by simulating and studying the actual stochastic epidemiological dynamics of multiple strains on concrete host contact networks. In that case, other topological properties of the networks, besides the degree distribution, such as clustering coefficient, could be examined. Moreover, to accurately represent real networks, degree correlations between nodes must be taken into account, a feature currently neglected in our study. Many real-world networks exhibit degree assortativity - the tendency for nodes with similar degrees to connect. As suggested by previous work [17], the effect of degree assortativity (or disassortativity) on long-term SIS disease dynamics could be substantial. Similarly, although we assumed a static network degree distribution, multi-strain propagation on dynamic host networks could be studied in the future, both for analytical and computational advances, along the same lines adopted here. Moreover, in this paper, we focused only on the special case where Λ depends on *µ* (Eq.2.16), ignoring the other components of the perturbations in Eq.2.13. This is, as motivated throughout the paper, the only perturbation that remains non-trivial when considered in isolation. The reason is that, unlike the other perturbations, it involves a global parameter that cannot be factorized, and therefore affects the structure of the dynamics and not only their timescale [11]. Nevertheless, if one wishes to explore the effect of the other perturbations, they should of course be included, as they are likely to generate very interesting selection patterns even for *N* = 2 in homogeneous populations [19].

Another central aspect that might be limiting the application of our findings is the quasi-neutrality assumption in the derivation of the formalism. Deviations from it, namely having strains that largely differ from each other won’t allow the connection between underlying host population contact network and macroscopic strain dynamics in the form of an emergent replicator, and may require a different approach.

### Implications for vaccination effects and multi-type disease control

The integration of network theory, epidemiology, and replicator dynamics provides a rigorous framework to study the global competition between pathogen strains and their impact on epidemic outcomes. In this setting, the replicator dynamics unraveled here for the network describes the selective success of different pathogen variants, driven by their relative fitness on one hand, and compounded by the heterogeneity in the underlying host contact network on the other.

By linking epidemiology with network theory and explicitly capturing the impact of network heterogeneity, our framework provides a tool to better understand how interacting strain frequencies evolve within structured populations, and are shaped by interventions. This approach could be especially important for extending the study of vaccination effects on networks [32] to polymorphic endemic systems or multi-type infectious agents.

Bacterial infectious diseases such as those caused by *Streptococcus pneumoniae* serotypes could be a possible venue of such model application, where the search for optimized universal as well as serotype-specific vaccines remains an area of active investigation [35]. Coupling such vaccination details with multi-strain dynamics on explicit host contact networks will provide an excellent venue for designing even more precise targeted immunization.

Finally, our study has implications not only for interacting multi-strain disease epidemiology and mathematical modeling, but more generally for the understanding of biodiversity and interacting phenomena propagation in realistically structured populations.

## Acknowledgements

This work was funded by the Portuguese Foundation for Science and Technology (FCT) via grant number 2022.03060.PTDC (Models4Invasion) and supported by the European Union (grant 101080528 - NOSEVAC).

## Code availability

Some codes related to the study are available at [GitHub](https://github.com/stenmadec/SIS_Host_Heterogeneity).

## Supplementary Information

## S1 Details on the explicit matrix of the replicator equation

Let us recall the replicator equation (2.12) used in this text:

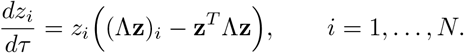

As proven in [21], the pairwise invasion fitness matrix between strains 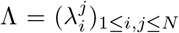 is given by (2.13). It depends on the global quantity *µ* (average ratio of single- to coinfection in the system) defined in (2.15) — which is extensively discussed in this text — and also on the five emergent network mean-field quantities Ξ_*s*_, *s* = 1, …, 5, defined explicitly as functions of the epidemiological parameters, network degree distribution, and endemic prevalence equilibrium:

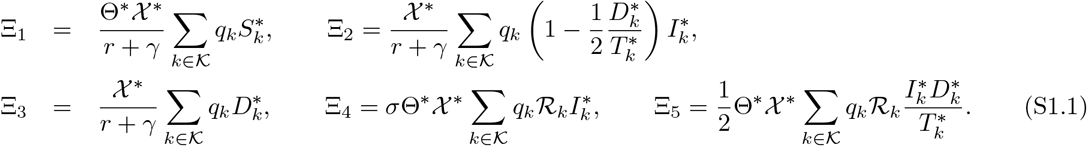

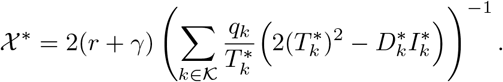

### S1.1 Explicit formulation for the special case of perturbation only in co-colonization interaction coefficients

In the special case we consider in the main text, we have only the trait 5 perturbation appearing in the pairwise invasion fitness. Let’s simplify the Ξ_5_ expression.

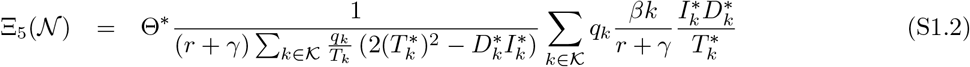

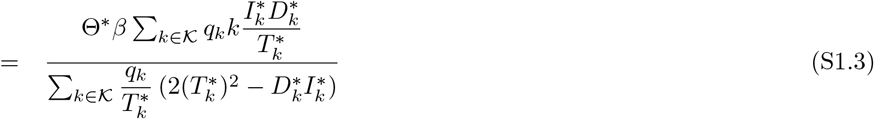

Recall that in the corresponding homogeneous system (e.g. [19]), the corresponding final weight for this perturbation is:

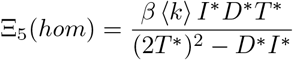

We have

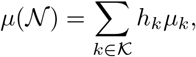

with 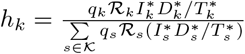, where 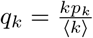.

Recall that

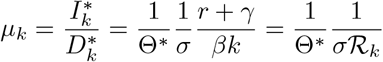

So comparing the two multi-strain systems (homogeneous population vs. network) will come down to comparing two replicators whose payoff matrices satisfy the following structure coefficient-wise:

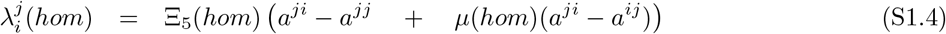

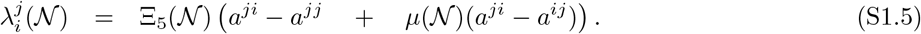

## S2 Additional numerical simulation results

**Figure S1:**
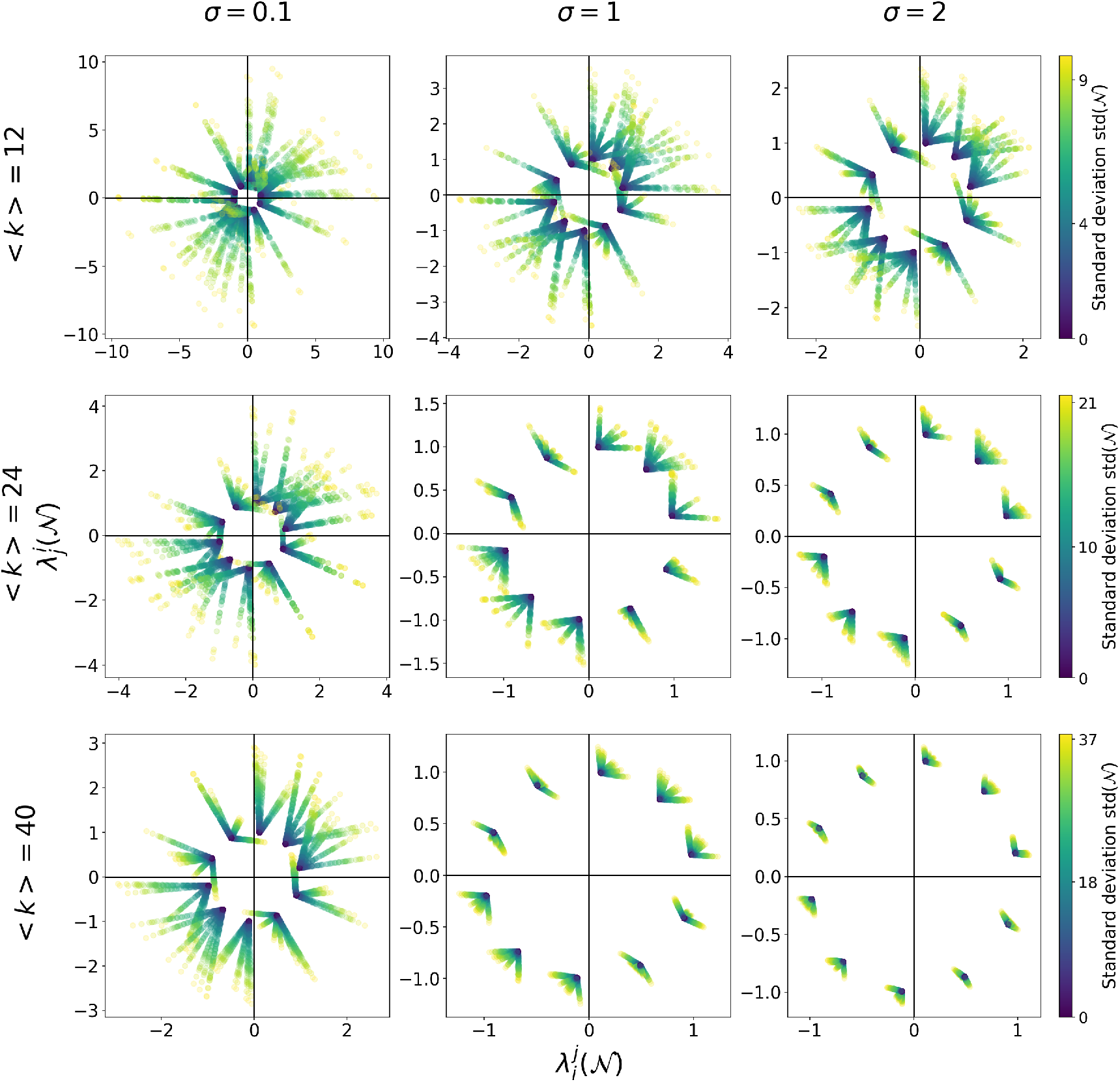
Effect of network heterogeneity on the pairwise invasion fitnesses of 2 strains 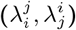 in the case of *N* = 2 for three different *R*_0_ (here changed via ⟨*k*⟩) and three different values of *σ*, under a different topology. All the networks are generated randomly within the *three-class* topology and have the same ⟨*k*⟩ in each row. Here N_*three*_(*s, p*) = {[⟨*k*⟩ −*s*, ⟨*k*⟩, ⟨*k*⟩ + *s*], [*p*, 1 −2*p, p*]}. We observe again lower sensitivity to network heterogeneity as transmission intensity and among-strain cooperation in coinfection increase.

**Figure S2:**
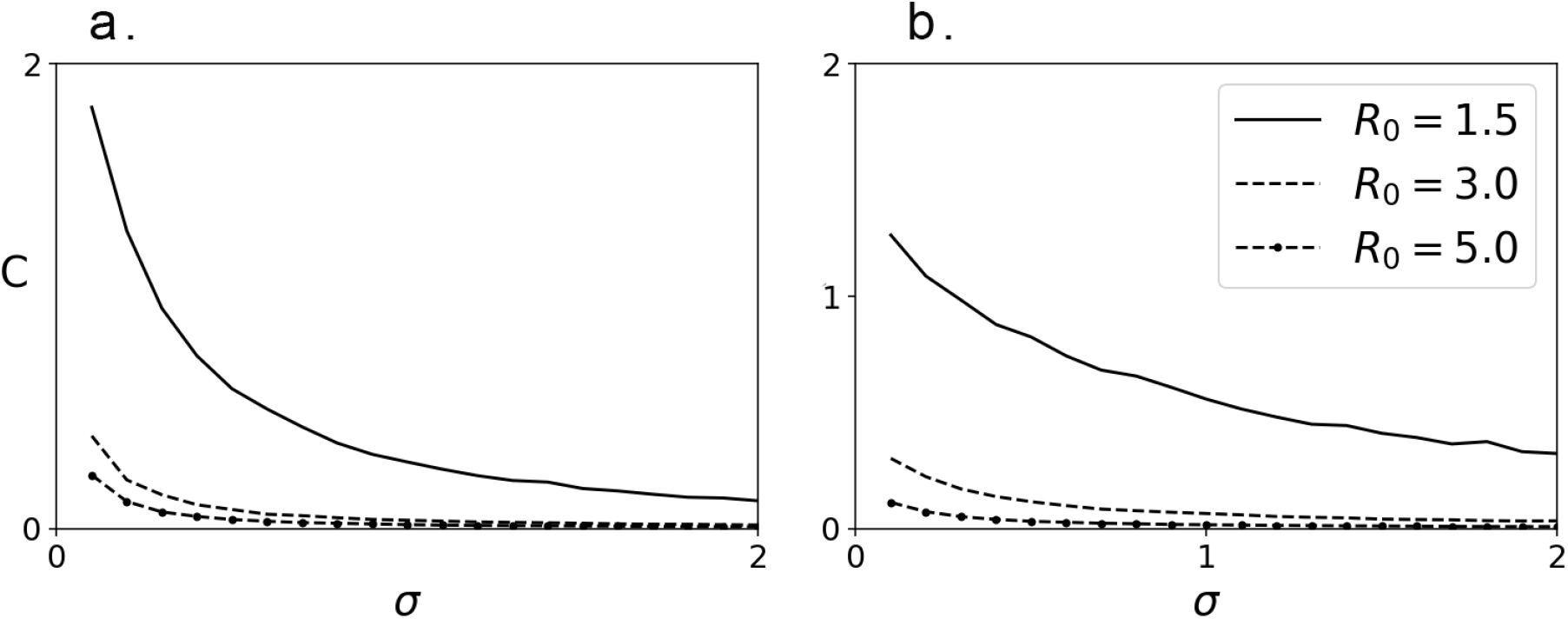
Sensitivity of replicator Λ matrix (for *N* = 2) to the network heterogeneity decreases with mean coinfection susceptibility factor *σ*. For each couple of fixed values *R*_0_ and *std*(N), we compute the Λ distance from homogeneity *d*(N, *A*) as in Eq. 4.1 (Fig. 3). We compute *d* for each of the six matrices *A* within the ten homogeneously equivalent classes and each network N, and regress all the obtained distances *d* vs. standard deviations *std*(N) of their corresponding networks, as in Eq. 4.2, saving the slopes as a measure of sensitivity to network heterogeneity. Stratifying them by *R*_0_ and *σ* we obtain *C*(*R*_0_, *σ* ), plotted here. We plot sensitivity to heterogeneity *C* (in the case of *N* = 2 strains) with respect to the epidemiological parameters *σ* and *R*_0_ for: **(a.)** fully random network topology and **(b.)** three-degree-class network topology, N_*three*_(*s, p*) = {[⟨*k*⟩ −*s*, ⟨*k*⟩, ⟨*k*⟩ + *s*], [*p*, 1 −2*p, p*]}. *C* is the least square regression coefficient computed from the distance from homogeneity of the pairwise fitnesses as a function of the standard deviation of the networks. In this example we have *N*_*k*_ = 3 values of *R*_0_ = ⟨*k*⟩×*ρ*_0_ and *N*_*σ*_ = 20 values of *σ* ∈ {*k/*10 for *k* = 1,*·*··, 20}. For each fixed *R*_0_ and *σ* we have *N*_*net*_ = 100 randomly generated networks *N* with the same ⟨*k*⟩ but different *std*(*N*) and we fix *N*_*λ*_ = 10 values of homogeneous 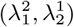 for which we generated *N*_*A*_ = 6 different *A*’s. Finally, for each of the *N*_*k*_ × *N*_*σ*_ = 60 couple (*R*_0_, *σ* ) we got *N*_*A*_ × *N*_*λ*_ × *N*_*net*_ = 6000 pair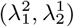. Each of these pairs is at a given distance *d*_*o*_ from the pair in the corresponding homogeneous network, yielding 6000 values of *d*_0_. We then plot *d*_0_ as a function of *std*(N), which leads to a cloud of points similar to figure 3.b, and we compute *C* = *cov*(*d, V* )*/V ar*(*V* ), the regression slope. Hence, for each of the 60 values of (*R*_0_, *σ* ), *C* is the summary of 6000 pairwise fitness.

**Figure S3:**
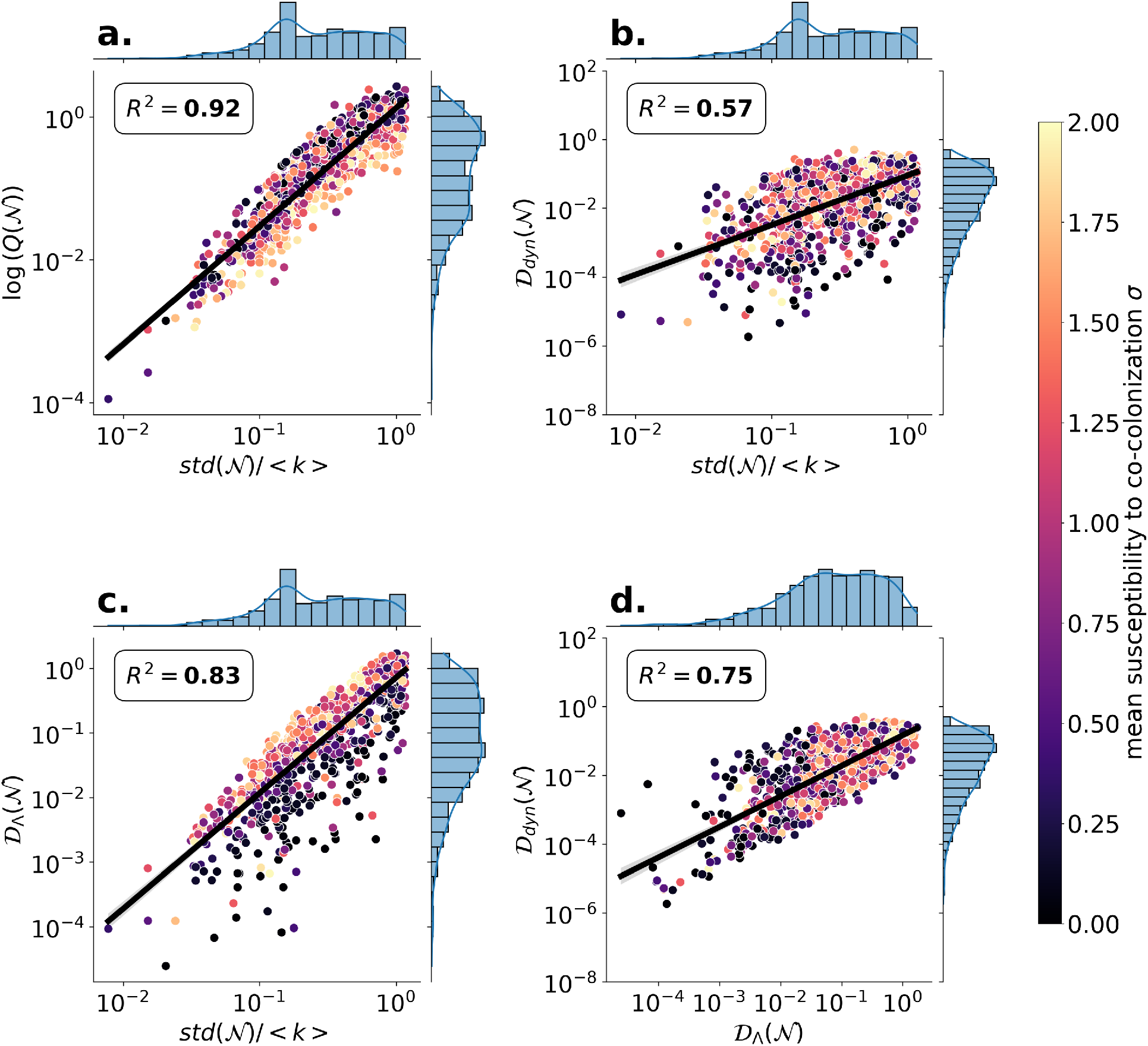
Stratification of simulations of Figure 6, by mean interaction in coinfection. We add the information of *σ* values to each simulated dynamics on a host contact network via a color gradient, reflecting values closer to 0 (competition) and above 1 (cooperation). Formal linear regression tests with COV and *σ* as independent variables were conducted, testing for their independent and interaction effects on the four *y* key observables depicted in a-d. **a**. <u>Increasing *σ* (cooperation) decreases the speed with a coefficient of -0.18</u>. However the value of sigma itself does not affect the relationship between network CV and speed of multi-strain dynamics (p-value for interaction =0.203). **b**. We find again no significant interaction between *σ* and COV, but here <u>both *σ* and COV independently increase the dynamic distance</u> of the multi-strain dynamics from the homogeneous counterpart. **c**. The <u>Λ distance increases significantly with *σ*</u>, with a coefficient of 0.32, and with the COV with a coefficient of 1.9. The latter suggests that in systems with more cooperation between strains, the distance between multi-strain dynamics on a network and in a homogeneous population is bigger. Interestingly, here we find also a relatively small but significant interaction between *σ* and COV (coefficient=-0.14, p-value=0.013), namely the sensitivity to the network CV decreases with sigma. This implies that when there is more cooperation between strains, the effect of network variance in degree distribution has a weaker effect on the payoff matrix of the corresponding replicator system for strain dynamics. This is in line with what we expect from the *N* = 2 case, when increasing cooperation was seen to affect less the lambda coefficients (Figure 4). **d**. We find that *σ* by itself has no significant effect, thus the level of cooperation or competition among strains in general has no significant average effect on dynamic distance when the lambda distance is zero, but in interaction with lambda distance it becomes weakly significant (coefficient 0.092, p-value =0.028). The positive relationship expected between Λ distance and dynamic distance in a replicator increases more in systems with more cooperation among strains. (Notice that in contrast to Figure 4,S1,S2, here, the baseline systems are not fixed, they are *σ* -dependent.)

**Figure S4:**
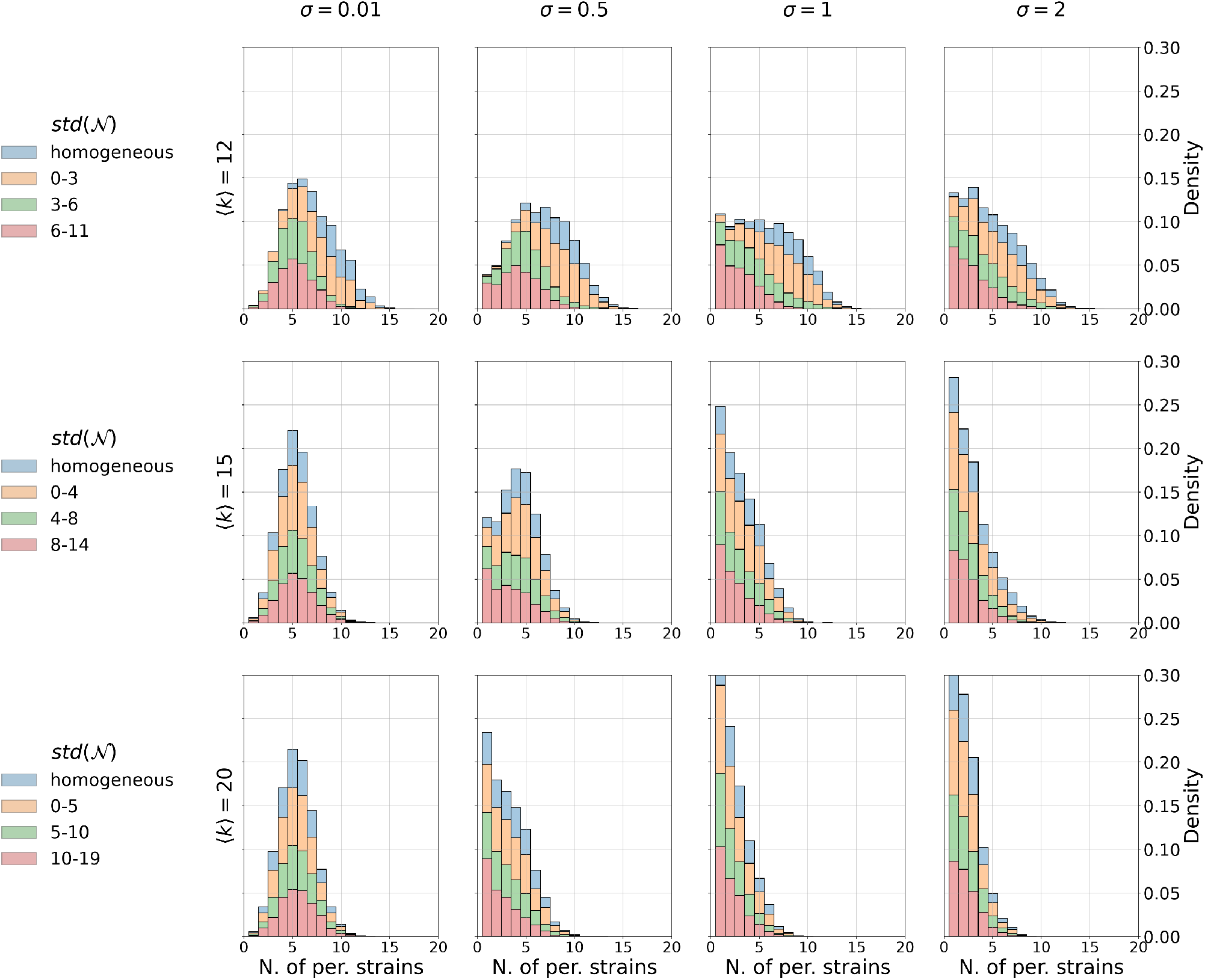
Persisting strains with respect to ⟨*k*⟩, *σ*, and *std*(N). Here *β/*(*r* + *γ*) = 0.1. For each value of ⟨*k*⟩ and *σ*, the system is split into four classes of *std* as shown in the legends. We compute 2500 replicator systems with *N* = 20 strains, using random *α*’s and a random network within the corresponding *std* range. The Λ matrices are renormalized by Λ_*norm*_ in order to have the same time scale *speed* (N) for each simulation. The dynamics are initialized at 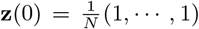 and evaluated at *T*_*max*_ = 5000. The histogram shows the number of persisting strains (above 0.1*/N* ) at *T*_*max*_. The complete histogram is reconstructed by computing the weight of each class from 10k random network launches using the same procedure.

**Figure S5:**
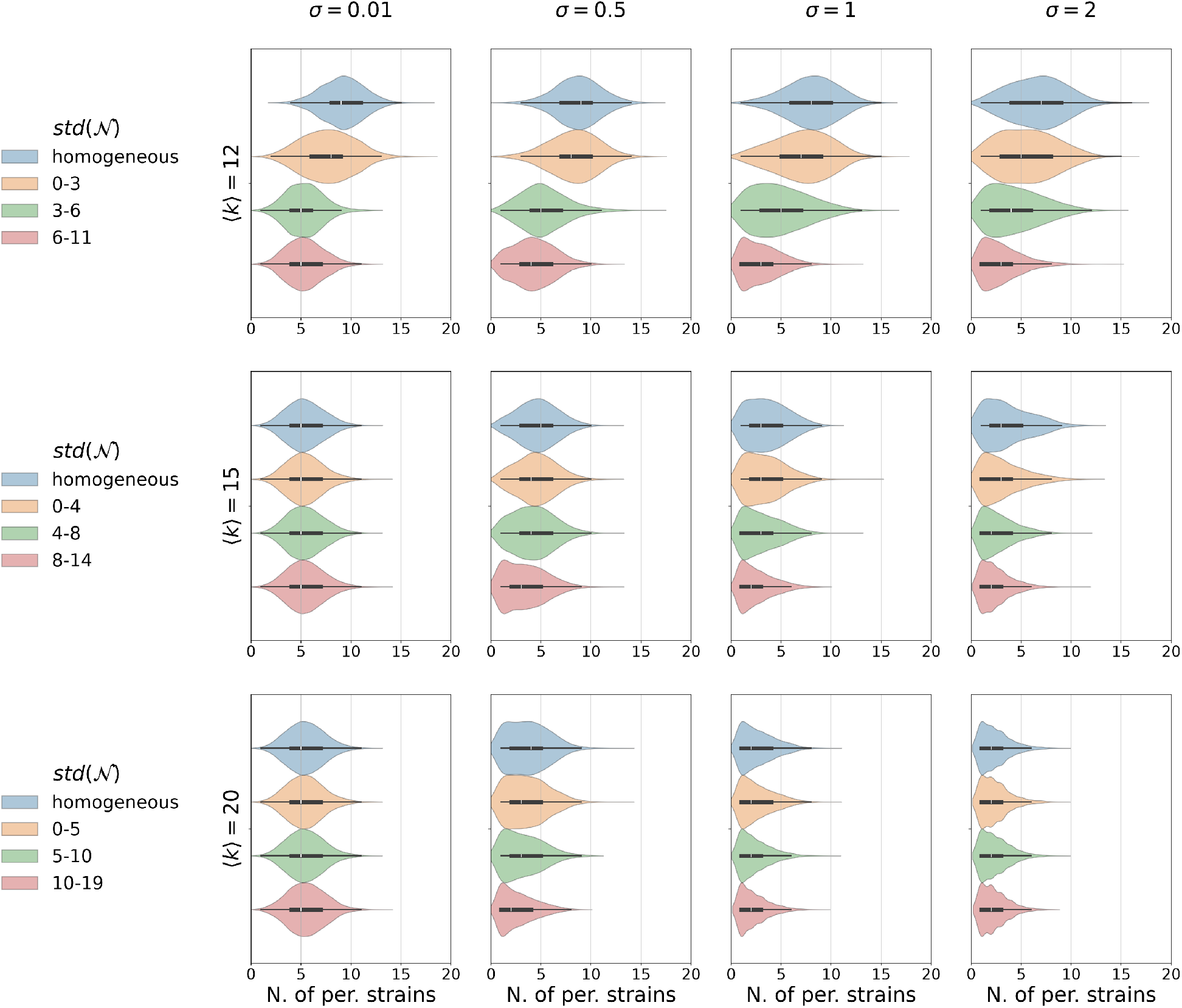
Number of persisting strains in different networks and parameter regimes - another summary representation of simulations in Figure S4. There is more sensitivity to the network heterogeneity for smaller transmission intensity and strain interactions in co-infection tending toward mean competition (*σ* smaller).

**Figure S6:**
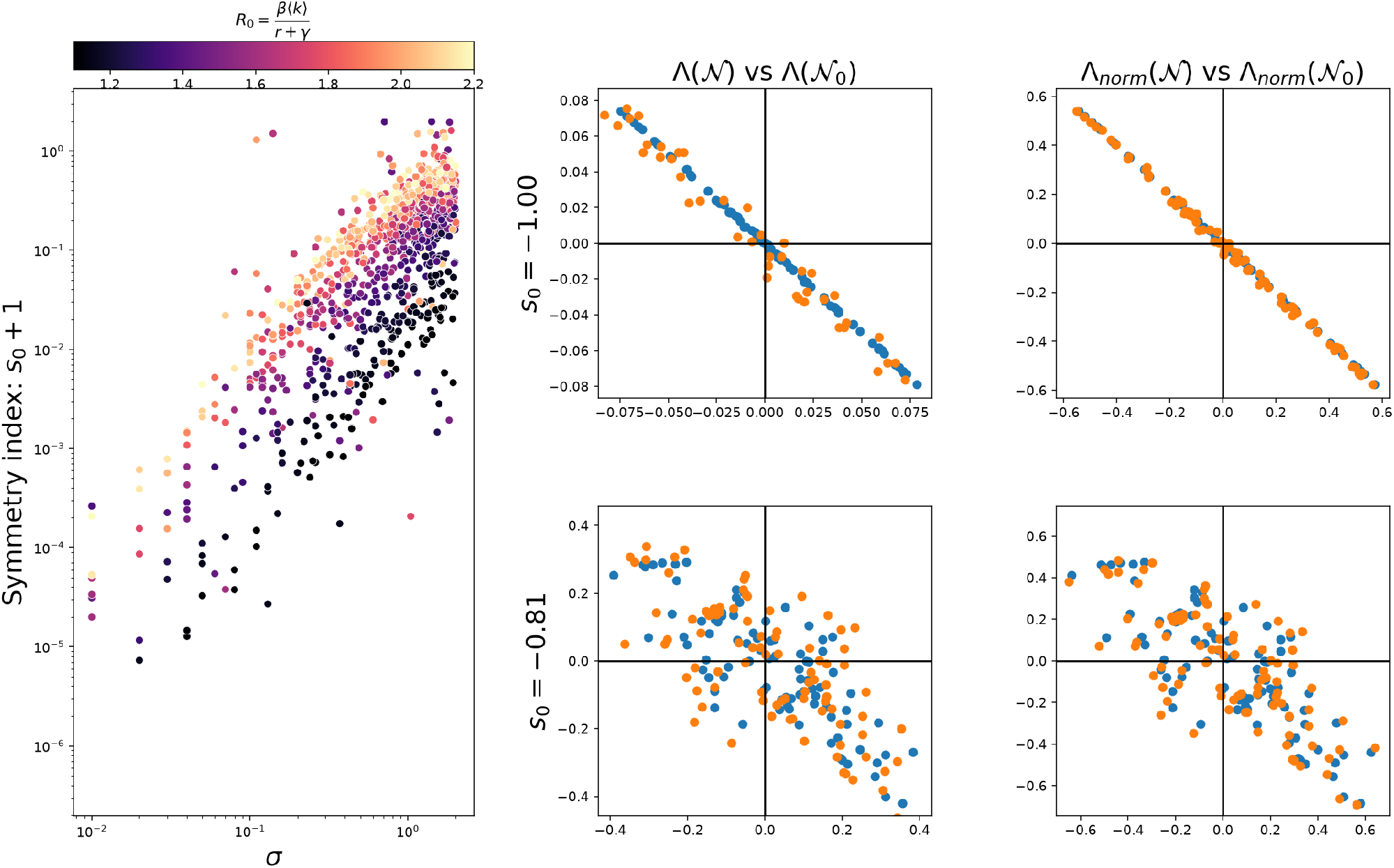
Illustration of Λ symmetry effect on the invasion fitness matrix comparison after normalisation. Left panel. Link between *σ* and *R*_0_ and the symmetry index of the baseline system Λ matrix 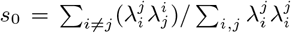. **Four right panels**. How symmetry affects Λ matrix renormalisation and network effects on multi-strain replicator dynamics. We illustrate for two different baseline systems: in the top row a system with higher anti-symmetry in the baseline Λ matrix, in the bottom row, a system with a lower anti-symmetry index in the baseline Λ matrix. (Blue : homogeneous system pairwise invasion fitness coefficients depicted as points on the plane, Orange, corresponding coefficients in the heterogeneous host contact network). We notice a relatively higher effect of the network, distance between blue and orange points, in the second case, where the baseline system is less anti-symmetric. These are the types of dynamics where the network effect will be felt more strongly and may induce bigger qualitative shifts in strain selection.

## Footnotes

1 We use the notation *β* for convenience, but here it refers to a probability of infection per contact, in contrast to the symbol *β* describing transmission rate in homogeneous mixing models [22].

2 Throughout the paper, we focus on finite networks, hence, the variance of the degree is always well-defined, and we do not deal with divergent variance [4].

3 Strictly speaking, this should be denoted by 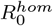 but for simplicity we will refer to it from now onwards as *R*_0_.

## Notes

https://github.com/stenmadec/SIS_Host_Heterogeneity

### Competing Interest Statement

The authors have declared no competing interest.

### Summary of Updates

We fixed some typos and equation formulations. We performed additional simulations. We clarified interpretation and the role of different parameters, in particular the role of coinfection susceptibility factor. We made some quantities explicit, including basic reproduction numbers in each class, the pairwise invasion fitness weights and their comparison with the homogeneous model. We improved interpretation of model results.

